# A Conserved Core Region of the Scaffold NEMO is Essential for Signal-induced Conformational Change and Liquid-liquid Phase Separation

**DOI:** 10.1101/2023.05.25.542299

**Authors:** Christopher J. DiRusso, Anthony M. DeMaria, Judy Wong, Jack J. Jordanides, Adrian Whitty, Karen N. Allen, Thomas D. Gilmore

**Author notes:** Corresponding authors: Thomas D. Gilmore, Biology Department, Boston University, 5 Cummington Mall, Boston, MA 02215, USA, Karen N. Allen, Department of Chemistry, Boston University, 590 Commonwealth Ave, Boston, MA 02215, USA. These authors contributed equally to this work (Christopher J. DiRusso, Anthony M. DeMaria). **Abbreviations:** The abbreviations used are: aa (amino acids), IFN (interferon), IKK (IκB kinase), IκB (inhibitor of κB binding), IVD (intervening domain), KBD (kinase binding domain), KO (knockout), LIR (LC3 Interacting Region), LLPS (liquid-liquid phase separation), LZ (leucine zipper), NEMO (NF-κB Essential Modulator), NF-κB (Nuclear Factor kappaB), OPTN (optineurin), SDS-PAGE (sodium dodecyl sulfate polyacrylamide gel electrophoresis), TNF (tumor necrosis factor), Ub (ubiquitin), UBAN ((Ubiquitin Binding in ABIN and NEMO domain), WT (wild-type), ZF (zinc finger).

## Abstract

Scaffold proteins help mediate interactions between protein partners, often to optimize intracellular signaling. Herein, we use comparative, biochemical, biophysical, molecular, and cellular approaches to investigate how the scaffold protein NEMO contributes to signaling in the NF-κB pathway. Comparison of NEMO and the related protein optineurin from a variety of evolutionarily distant organisms revealed that a central region of NEMO, called the Intervening Domain (IVD), is conserved between NEMO and optineurin. Previous studies have shown that this central core region of the IVD is required for cytokine-induced activation of IκB kinase (IKK). We show that the analogous region of optineurin can functionally replace the core region of the NEMO IVD. We also show that an intact IVD is required for the formation of disulfide-bonded dimers of NEMO. Moreover, inactivating mutations in this core region abrogate the ability of NEMO to form ubiquitin-induced liquid-liquid phase separation droplets in vitro and signal-induced puncta in vivo. Thermal and chemical denaturation studies of truncated NEMO variants indicate that the IVD, while not intrinsically destabilizing, can reduce the stability of surrounding regions of NEMO, due to the conflicting structural demands imparted on this region by flanking upstream and downstream domains. This conformational strain in the IVD mediates allosteric communication between N- and C-terminal regions of NEMO. Overall, these results support a model in which the IVD of NEMO participates in signal-induced activation of the IKK/NF-κB pathway by acting as a mediator of conformational changes in NEMO.

## Introduction

Intracellular processes require the precise coordination of molecular components for optimal output. In many cases, scaffold proteins facilitate these intracellular processes by acting as docking hubs for interacting partners in a given signaling pathway (1). However, scaffold proteins are not simply inert frameworks for such coordinated events. That is, scaffold proteins can themselves contribute to these multicomponent molecular processes by undergoing controlled conformational changes that bring multiple components into the proper configuration for downstream signaling.

One such multicomponent process is the transcription factor NF-κB pathway, which regulates myriad cellular and organismal processes including immunity, development, and cell survival (2). In most resting cells, NF-κB dimers are sequestered in the cytoplasm in a complex with inhibitor of NF-κB (IκB) proteins, which are substrates for regulated phosphorylation by the IκB kinase (IKK). The canonical IKK complex is composed of the kinases IKKα and IKKβ, and the scaffold NEMO (also called IKKγ) (3). In canonical NF-κB signaling, activated IKKβ carries out the phosphorylation of IκB (primarily IκBα), which is then followed by degradation of IκB, and the liberation of NF-κB for translocation to the nucleus and activation of NF-κB target genes (3, 4). Knockouts and mutations of NEMO abolish the ability of cells to respond to canonical NF-κB pathway activators, such as tumor necrosis factor alpha (TNFα) and interferon beta (IFNβ) (5, 6). NEMO is an essential scaffold for the IKKβ-dependent phosphorylation of IκBα, but the mechanism by which NEMO exerts its scaffolding activity is not completely understood.

Human NEMO is a 419 amino acid protein with several subdomains that contribute to its scaffolding function and hence NF-κB signaling. As shown in Fig. 1A, these domains include a primary N-terminal binding domain for IκB kinase beta (IKKβ) (KBD, ∼aa 44-111), a ubiquitin-binding domain (UBAN, ∼aa 286-321), and a C-terminal zinc finger domain (ZF, ∼aa 389-419), which has been implicated in IκBα binding.(7–10). One region of NEMO (∼aa 110-195, called the Intervening Domain [IVD]) is not known to bind any other proteins/ligands, and its precise function is not known. Nevertheless, some mutations in conserved core sequences of the IVD abolish the ability of NEMO to support cytokine-induced activation of IKK (5, 11). Although the overall structure of NEMO is likely to be a series of coiled-coil domains, the X-ray crystallographic structure of the IVD has yet to be determined.

**Figure 1.**
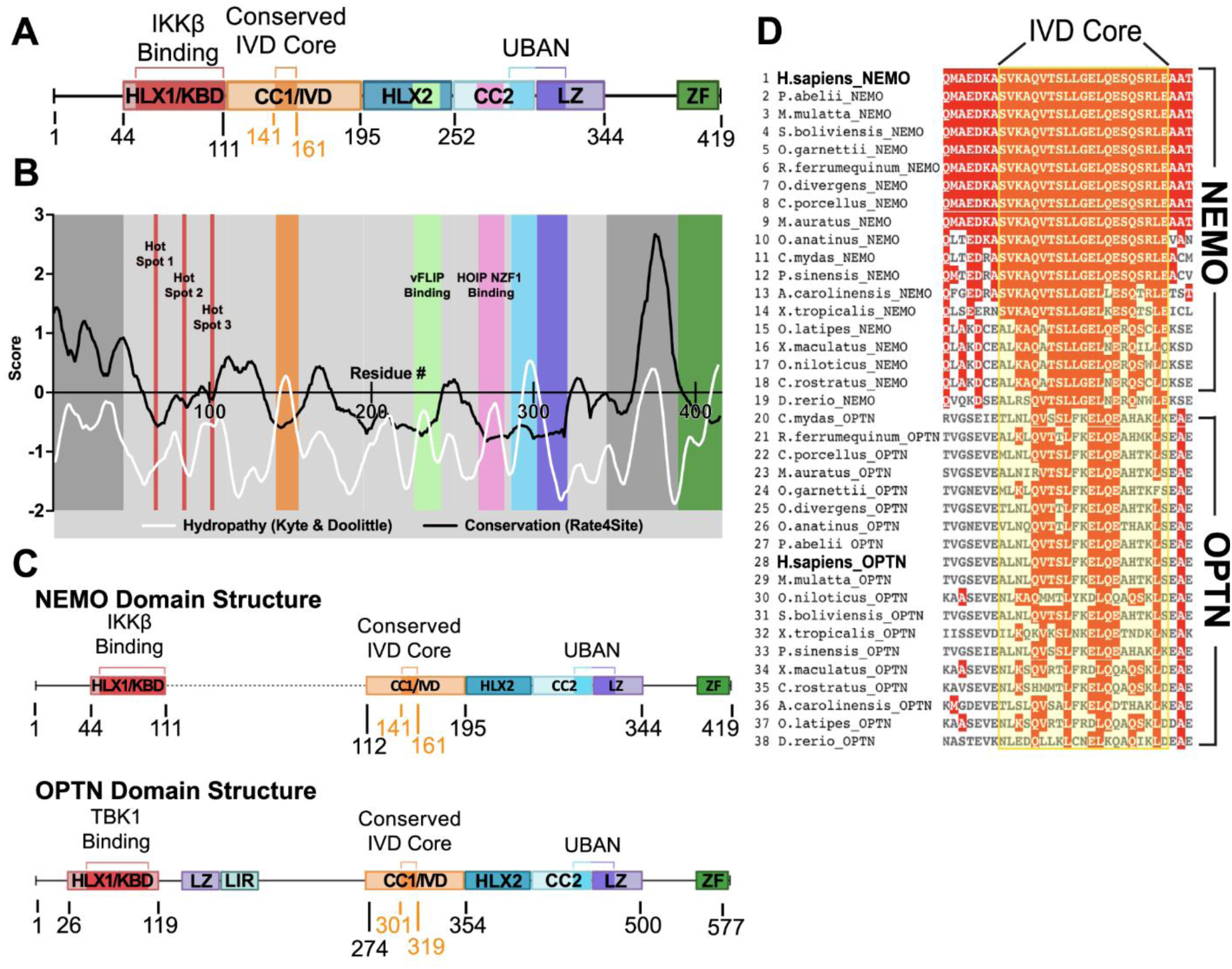
Evolutionary analysis and comparison of NEMO and OPTN proteins. (A) NEMO domain structure, with colored areas corresponding to the functional domains of NEMO and proposed unstructured regions represented by black lines. HLX1/KBD, Helix 1/Kinase Binding Domain; CC1/IVD, Coiled Coil 1/Intervening Domain; HLX2, Helix 2; CC2, Coiled Coil 2; LZ, Leucine Zipper; UBAN, Ubiquitin Binding in ABIN and NEMO domain; ZF, Zinc Finger. (B) Plot of the sequence conservation obtained using Rate4Site and the residue hydropathy calculated using the Kyte & Doolittle scale for human NEMO, represented by a moving average. The red bars denote the location of the hot spot regions of the NEMO-IKKβ binding interaction (Fig. S1). (C) Structural comparison of NEMO and OPTN protein domain structures, with the black dotted line indicating a gap in the alignment. LIR, LC3 Interacting Region. (D) Multiple sequence alignment for the IVD core region of NEMO and OPTN, from a broad spread of vertebrate proteins. Sequences were aligned with Clustal Omega (EMBL-EBI) using WT human NEMO as a reference sequence. The sequence alignment was visualized and colored according to residue conservation using MView (EMBL-EBI) (56).

Several studies have identified NEMO as a dynamic regulator of canonical NF-κB activation, primarily through ligand and ubiquitin-induced conformational plasticity (12–16). For example, Shaffer et al. showed that NEMO becomes more compact upon binding to IKKβ (13), and Hauenstein et al. (12) and Catici et al. (16, 17) demonstrated that binding of linear ubiquitin leads to a variety of distinct conformational states of NEMO. Finally, two recent reports showed NEMO undergoes signal-induced liquid-liquid phase separation (LLPS) upon binding ubiquitin (18, 19).

Mutations in NEMO that reduce or abolish its ability to support downstream activation of NF-κB are the cause of several human immunodeficiency diseases (5). In many cases, these mutations interfere with the ability of NEMO to interact with its key protein partners (8, 9, 20, 21). Nevertheless, several human disease mutations located in the IVD are not known to affect NEMO-dependent protein-protein interactions, even though they reduce the ability of NEMO to support cytokine-induced activation of NF-κB (5). Moreover, we have previously shown that a substitution at a highly conserved region (aa 145-153, replaced by a Ser-Gly repeat; mutant 9SG) of the IVD abolishes the ability of NEMO to support TNFα-induced phosphorylation of IκBα, although this mutation does not block the interaction of NEMO with IKKβ (11, 13).

In this study, we have used comparative, molecular, biochemical, biophysical, and cellular approaches in an effort to understand the role of the IVD in conformational changes in NEMO that are required for activation of IKK. Our results support a model in which the IVD is a sequence-dependent driver of conformational changes and LLPS that are required for activation of IKK/NF-κB. As such, these findings suggest that mutations in the IVD affect the ability of NEMO to undergo stimulus-induced conformational change and can be classified as a new type of scaffolding mutation leading to disease. Our model provides insights for therapeutic design for disease treatment.

## Results

### Comparative sequence analyses of the IVD region of NEMO suggest functional importance

Shaffer et al.(13) highlighted the highly conserved core region of the IVD as critical for NEMO function and proposed a role for IVD-mediated conformational change in the downstream signal propagation of the NF-κB pathway. To provide additional insight into the structure/function relationship in the IVD, we determined how the evolutionary rate of the IVD compared to other structurally characterized domains of NEMO that participate in protein-protein interactions. We used the tool Rate4Site (22) to construct a phylogenetic tree from a multiple-sequence alignment, using the neighbor-joining algorithm, which estimates the maximum likelihood evolutionary rate for each residue position. The rates are normalized around zero with a standard deviation of one, with positive scores representing higher and negative scores representing lower rates of evolution. We used 500 NEMO sequences from vertebrate species (sequence identity of 64-100%) to perform this analysis using a four-period moving average to better uncover trends of functional importance in NEMO (Fig. 1B). The analysis shows a region of low evolutionary rate that coincides with the 44-111 KBD region of NEMO. The affinity of the partners in this protein-protein binding interface is dependent on three “hot spots” which were experimentally validated and independently identified by computational structural mapping (23). These hot spots align well with the sites of lowest evolutionary rate in the KBD region (Fig. S1). In addition, the HLX-2 (Helix 2) domain, the CC2 domain, and the UBAN region that spans the CC2 and LZ domains, which contain defined segments that participate in protein-protein interactions (24), all show low rates of evolutionary change in the expected regions. Therefore, the low evolutionary rate of the IVD (Fig. 1B) is similar to regions of NEMO that are known to be of functional importance. The evolutionary rate within the IVD is lowest for residues 141-161, suggesting a critical role for this core region. In contrast, the N-terminal disordered region of NEMO comprising residues 1-44 and the C-terminal disordered region (residues ∼345-388) have high rates of evolutionary divergence, suggesting they do not have functional roles that require sequence conservation.

As a second method of identifying key regions of NEMO, we performed a hydrophobicity analysis, using the Kyte & Doolittle scale to score the hydrophobicity of each residue subjected to a moving average. NEMO is primarily an extended coiled-coil protein with high solvent exposure, and thus displays a low average hydrophobicity. Therefore, especially hydrophobic regions are likely either to take part in protein-protein interactions or to comprise folded domains large enough to have a hydrophobic core. Indeed, the protein-interacting regions within the HLX-2, UBAN, and CC2 domains all display spikes of increased local hydrophobicity (Fig. 1B). Thus, there is a general correlation between low evolutionary rate and high hydrophobicity in these domains with known functional roles. Similarly, for the IVD, an area of high hydrophobicity overlaps with the area of low evolutionary rate, suggesting that the IVD is involved in protein-protein interactions and/or forms a compact folded domain (Fig. 1B).

The evolutionary rate analysis can be further used to score the order/disorder of a region and its propensity for forming protein-protein interactions by computing evolutionary ratios. Dubreuil et al. (25) determined that the average evolutionary rate of a disordered region divided by the average evolutionary rate of residues in the ordered domains of proteins from a large dataset of proteins of known structure yields a ratio of ∼ 2. For NEMO, we find ratios greater than 3 for both the N-terminal and C-terminal disordered regions, as expected, but a ratio of 1.4 for the central core of the IVD. The use of this ratio as a metric suggests that the IVD is not a fully disordered region, but it is not as ordered as other known structured domains of NEMO such as the HLX-2, CC2 and LZ (with ratios of 0.52, 0.81 and 1.1, respectively).

To identify possible homologs of NEMO, we generated a sequence similarity network (SSN) (Fig. S2) using human NEMO as a query. All proteins in the NEMO SSN cluster had 40-99% identity to human NEMO and appeared to be NEMO orthologs. A group of proteins with 22-31% sequence identity with human NEMO formed a second isofunctional cluster and were uniquely annotated as optineurin (OPTN), a protein whose sequence similarity to NEMO has been recognized previously (26).

Alignment of the complete sequences of 38 vertebrate NEMO and OPTN proteins (Fig. S3) showed that the domain organization of OPTN and NEMO are similar except for the presence of additional central LZ and LIR domains in OPTN (Fig. 1C). In particular, both NEMO and OPTN have extensive sequence similarity in the core IVD sequence (Fig. 1D).

### Substitution of the core sequence of the NEMO IVD with analogous sequences from OPTN does not affect the ability of NEMO to support TNFα-induced activation of IKKβ

Based on the sequence conservation of the core region of the IVD in both NEMO and OPTN proteins, we sought to determine whether these core regions were functionally similar. To do this, we created relevant mutations in the core region of NEMO, subcloned them into FLAG-tagged expression plasmids (Fig. 2A), and transfected the plasmids into 293T NEMO knockout cells (11). To assess the function of NEMO mutants, transfected cells were treated with TNFα, and TNFα-induced phosphorylation of IκBα was compared to empty vector-transfected cells (negative control) and to wild-type NEMO-transfected cells (positive control).

**Figure 2.**
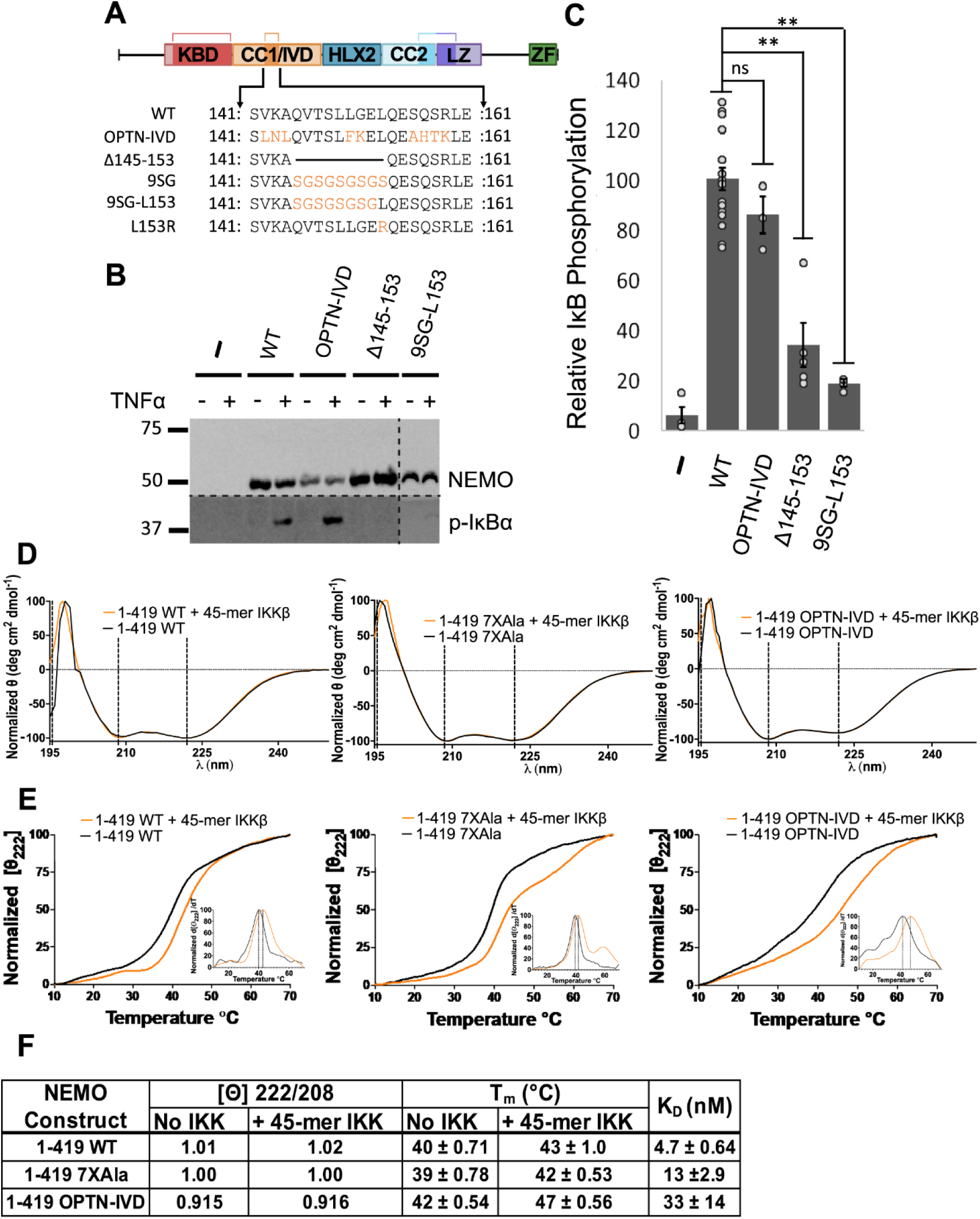
Effect of mutations in the core region of the IVD on the in vivo function and structural integrity of NEMO. (A) IVD mutants used in this study. Light blue residues indicate conservative amino acid substitutions, while orange residues indicate non-conservative substitutions. (B) Anti-NEMO (top) and anti-phospho-IκBα (bottom) Western blots of 293T NEMO KO cells transfected with the indicated FLAG-tagged NEMO expression vectors or an empty vector plasmid (-). Cells were treated with TNFα (final concentration 20 ng/ml) for 10 min prior to lysis (+) or left untreated (-) as indicated. Molecular weight markers (in kDa) are indicated to the left. (C) Quantitation of IκBα phosphorylation from at least three experiments per transfection condition. The amount of phosphorylation of IκBα is expressed as the ratio of phospho-IκBα to NEMO relative to average ratio of wild-type NEMO transfected cells (100). Shown are the means ± SEM; ns, not significant; *, p<0.05 by t-test assuming equal or unequal variance as appropriate; n= ≥ three individual experiments. Dots indicate values of individual experiments. (D) The CD spectra of indicated NEMO constructs determined at 25 °C at 10 µM, with and without equimolar equivalents of a 45-mer peptide of IKKβ, incubated for 1 h at room temperature. (E) CD-monitored thermal denaturation of NEMO constructs by measuring the loss in secondary structure as an increase in the 222 nm signal, ± equimolar equivalents of a 45-mer peptide of IKKβ, incubated for 1 h at room temperature. Insets show the peak of the first derivative as the melting temperature (T_m_). (F) Table of key CD results. The ellipticity ratios of the constructs is indicated as the ratio of the signal at 222 nm to 208 nm with and without the IKKβ peptide in (B), T_m_ with and without IKKβ peptide in (D), and K_D_ values for the binding affinity of the constructs for a FITC labelled 45-mer peptide of IKKβ determined using a fluorescence anisotropy assay (and see Fig. S5).

We first tested whether replacement of the most conserved region of NEMO’s IVD (aa 142-159) with the analogous region of OPTN (aa 302-319) affected NEMO’s signaling function in vivo. This segment of OPTN contains nine aa differences from the analogous region in NEMO, of which seven are conservative changes (V142L, K143N, L150F, S156A, Q157H, S158T, R159K) and two are not conservative (A144L, G151K) (Fig. 2A). The substitution mutant OPTN-IVD-NEMO showed no significant difference from wild-type NEMO in supporting TNFα-induced phosphorylation of IκBα (Figs. 2B, C). Thus, the conserved core IVD sequences of OPTN can functionally substitute for the core region of NEMO. In contrast, a NEMO mutant with a deletion of aa 145-153 (Δ145-153) showed a significant reduction in TNFα-induced phosphorylation of IκBα.

Since L153 is one of the most conserved residues (by sequence alignment, Fig. 1D) and the L153R mutation is detrimental to NEMO function (6, 11), we also tested if adding back only L153 would restore function to the inactive 9SG NEMO mutant, which has a repeating Ser-Gly replacement from aa 145-153 (11, 13) (Fig. 2A). This new mutant (9SG-L153) showed no significant ability to support phosphorylation of IκBα, confirming that the functional importance of the highly conserved core region of NEMO does not rely simply on L153. Taken together, these results confirm our comparative analysis showing that the core region of the IVD is important for NEMO function and that the analogous region of OPTN can functionally replace this core region of NEMO even though their sequences are not precisely conserved.

We also determined whether replacement of the core region of NEMO’s IVD with the corresponding segment of OPTN affected the ability of NEMO to bind IKKβ in vitro, using our previously established fluorescence anisotropy binding assay that uses an FITC-labeled 45-mer peptide derived from the C-terminal region of IKKβ (13, 23). For these assays, as well as the CD measurements described below, we used a form of NEMO in which seven Cys residues have been substituted with Ala (7XAla-NEMO, Fig. S4), because we have previously shown that this variant of NEMO is less susceptible to disulfide-bonded aggregate formation when expressed in bacteria and yet retains WT NEMO activity in human cells (27, 28). Full-length NEMO proteins (aa 1-1419), including WT NEMO, 7XAla-NEMO, and 9SG-NEMO, bind the 45-mer peptide derived from IKKβ with affinities ranging from 2-10 nM (23, 28). Using the same method, we found that OPTN-IVD-NEMO bound to the 45-mer peptide with K_D_ 33 ± 14 nM, which is slightly higher than the values seen for WT (4.7 ± 0.64 nM) or 7XAla -NEMO (13 ± 2.9 nM) (Figs. 2F, S5). Nevertheless, all three NEMO proteins fully support TNFα-induced activation of IKKβ in vivo.

To compare the structures of NEMO and OPTN-IVD-NEMO, we performed circular dichroism (CD) on bacterially expressed NEMO proteins in the presence and absence of bound IKKβ peptide (Fig. 2D). By CD, full-length WT NEMO and 7XAla-NEMO displayed nearly identical secondary structures strongly dominated by alpha helices, with a ratio of ellipticity at 222/208 nm of 1.02 and 1.00, respectively, indicating a high coiled-coil content. For OPTN-IVD-NEMO, we observed a similar α-helical structure with a decrease in the ellipticity ratio to 0.91, suggesting a small reduction in the coiled-coil content following the insertion of the IVD from OPTN into NEMO. We also compared the structural stability of the NEMO variants by determining their melting temperatures. Melting was measured by monitoring the change in the 222 nm peak in the CD spectrum as the temperature was ramped from 10 to 70 °C. OPTN-IVD NEMO was found to have a melting temperature of T_m_ = 42 ± 0.54 °C, while 7XAla and WT NEMO had T_m_ values of 39 ± 0.78 °C and 40 ± 0.71 °C, respectively (Fig. 2E) Thus, the OPTN-IVD substitution did not reduce the stability of the full-length NEMO protein. Moreover, inclusion of an equimolar concentration of the IKKβ 45-mer peptide caused an increase in T_m_ of 3 °C for WT NEMO and 7XAla-NEMO and of 5 °C for OPTN-IVD-NEMO. Overall, these results indicate that substitution of the OPTN-IVD in NEMO does not substantially alter full-length NEMO’s secondary structure, stability, or interaction with IKKβ.

### Disruption of the IVD impairs disulfide-bonded dimer formation of NEMO in vivo

Although the NEMO dimer is natively noncovalent in the reducing environment of the cell (28), certain cysteine residues, such as Cys54, are positioned at the dimer interface such that inter-chain disulfide bonding can occur if the protein is subjected to oxidizing conditions (29). We have previously shown that NEMO can form disulfide-bonded dimers in cells either when lysates are made in the absence of reducing agent or following treatment of cells with hydrogen peroxide (27, 28). To determine whether IVD mutations affect disulfide-bonded dimerization of NEMO, we first expressed FLAG-tagged versions of full-length WT, 9SG, and OPTN-IVD-NEMO proteins in 293T NEMO knockout cells. Cells were then lysed with a non-ionic detergent buffer and the lysates were analyzed by non-reducing (i.e., no β-mercaptoethanol or boiling) SDS-PAGE. We observed that WT NEMO migrated as two bands, with one band at the position expected for a NEMO monomer and a second higher band at approximately double the molecular weight, i.e., corresponding to dimeric NEMO. The active OPTN-IVD NEMO mutant also showed this two-band pattern and had a dimer/monomer ratio that was similar to that seen with WT NEMO. In contrast, no higher molecular-weight band was seen with 9SG NEMO (Fig. 3A), even though there are no Cys residue substitution mutations in 9SG. On fully denaturing SDS-PAGE bacterially expressed versions of these three proteins co-migrated as single bands (see below, Fig. 4D).

**Figure 3.**
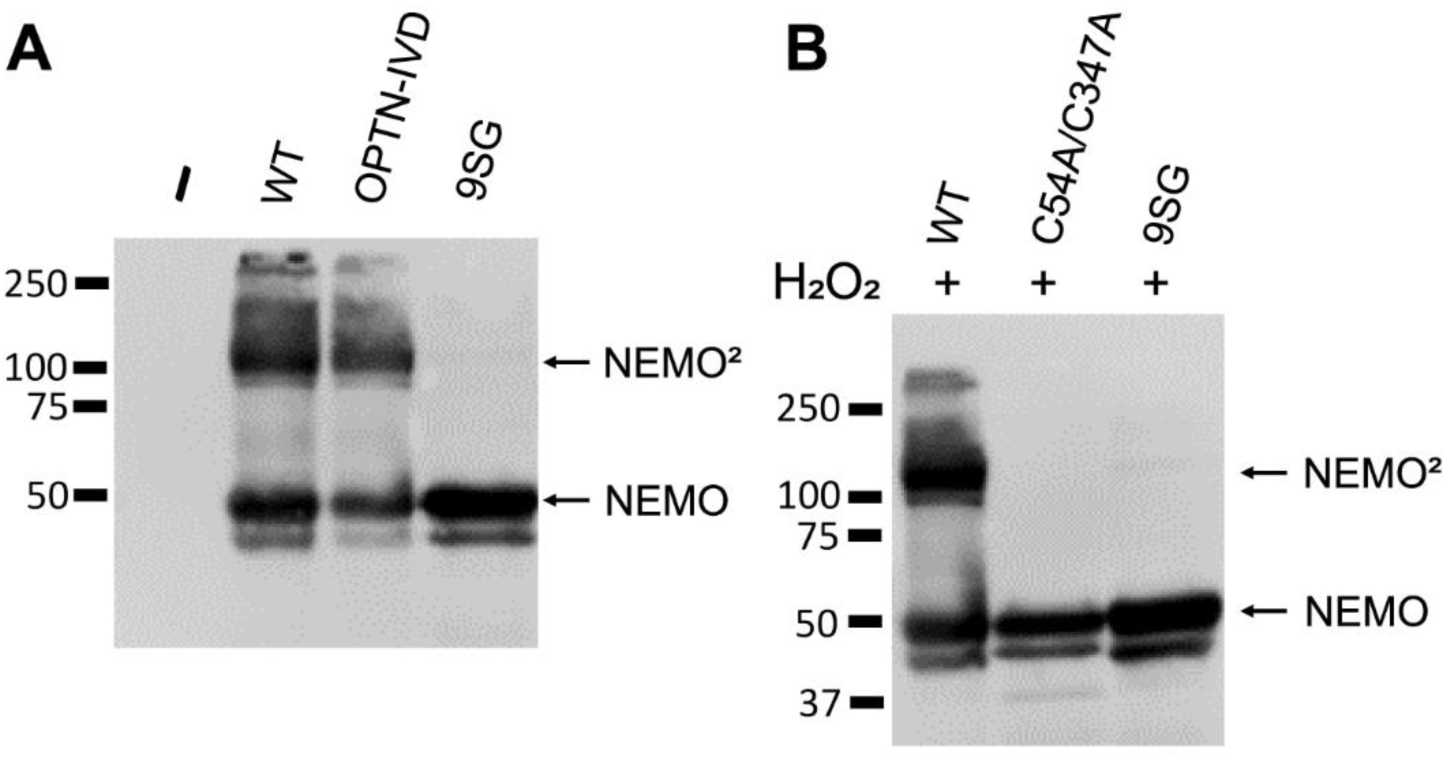
Mutations in a core region of the NEMO IVD affect its conformation and ability to form dimers. (A) Anti-NEMO Western blot of 293T NEMO KO cells transfected with FLAG-tagged expression vectors containing the indicated NEMO proteins or an empty vector (-). Samples were prepared in SDS sample buffer lacking β-mercaptoethanol and were not boiled prior to SDS-PAGE separation. NEMO, monomer, NEMO^2^, NEMO dimer. (B) Anti-NEMO Western blot of 293T NEMO KO cells transfected as in (A). Cells were treated with 50 µM hydrogen peroxide for 10 min prior to lysis. Samples were boiled in SDS-sample buffer lacking β-mercaptoethanol prior to separation by SDS-PAGE.

**Figure 4.**
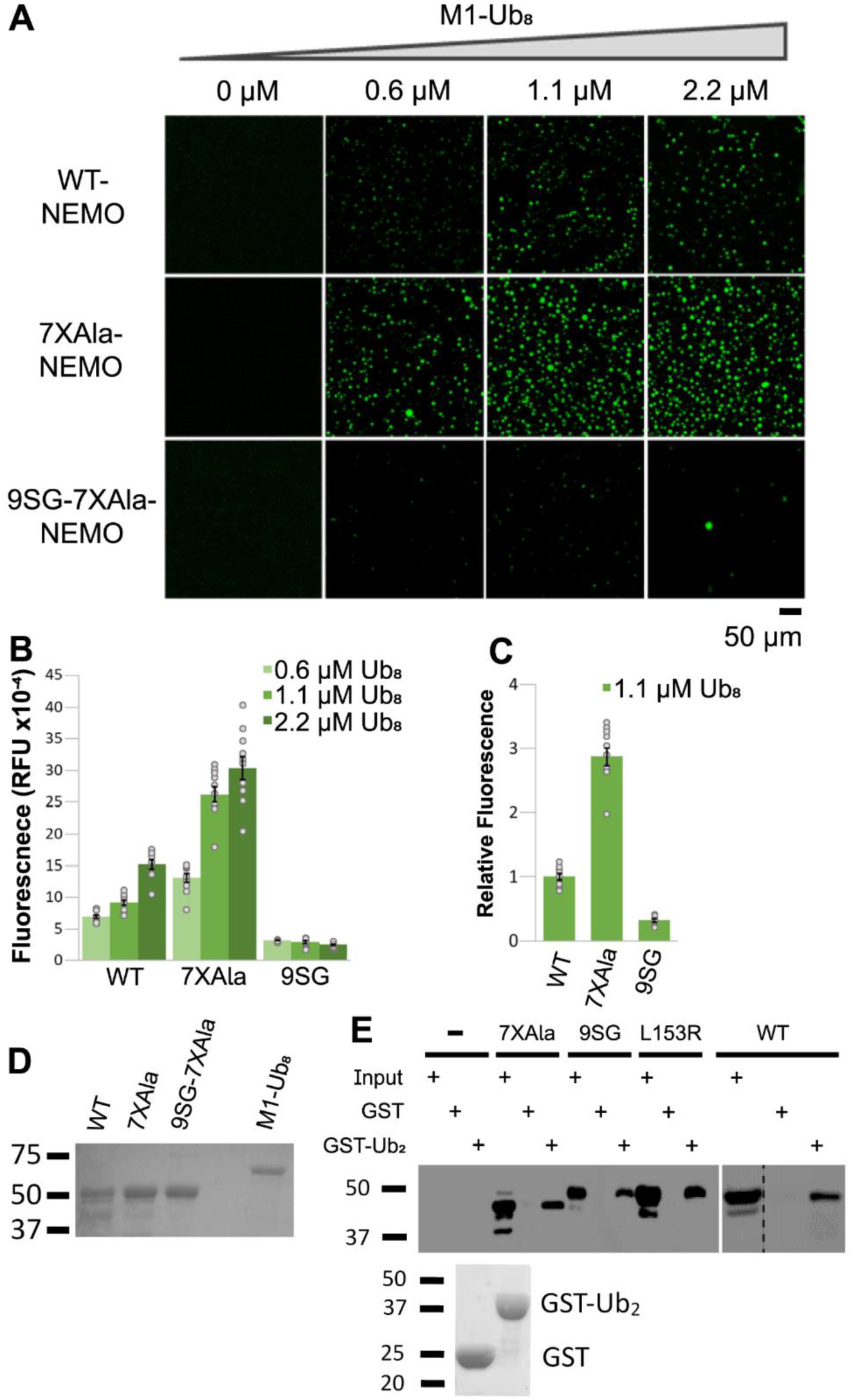
Mutation of the core region of the NEMO IVD abolishes its ability to undergo liquid-liquid phase separation. (A) Fluorescent images of liquid droplets of the indicated NEMO proteins induced by M1-Ub_8_ at different concentrations. Liquid droplets formed after mixing 6 µM of the indicated NEMO proteins with the indicated concentrations of M1-Ub_8_ for 45 min at 37°C. Scale bar, 50 µm. (B) Quantitation of fluorescence intensity of the liquid droplets in (A). Shown are the means ± SEM, n=10 areas, dots are the fluorescence intensities of individual areas. (C) Comparison of fluorescence at 1.1 µM Ub-8 for the indicated proteins. Values were normalized to the average intensity of WT NEMO (1.0). Shown are the means ± SEM, n=10 areas, dots are the fluorescence intensities of individual areas. (D) Coomassie blue-stained gel of the bacterially expressed proteins used in (A). Samples were prepared in SDS sample buffer containing β-mercaptoethanol and were boiled prior to SDS-PAGE. (E) Pull-down assay with GST and GST-Ub_2_ of the indicated NEMO proteins. The top panel shows an anti-FLAG Western blot of GST or GST-Ub_2_ pull-downs of lysates from 293T NEMO KO cells transfected with FLAG-tagged expression vectors containing the indicated NEMO proteins or an empty vector (-). Input lanes (I) contain 0.5% of the cell lysate used in each pulldown. The bottom panel is a Coomassie blue-stained gel of 1% of the GST and GST-Ub_2_ proteins used in the pulldowns.

As a second measure of oxidative dimer formation, we assessed the ability of the NEMO 9SG mutant to form disulfide-bonded dimers in cells following hydrogen peroxide treatment. We have previously shown this disulfide-bonded dimerization of NEMO involves Cys54 and Cys 347 (27). Therefore, we transfected 293T NEMO KO cells with FLAG-tagged versions of full-length WT, 9SG (WT background), and C54/347A NEMO proteins, treated the cells with hydrogen peroxide, and then subjected the lysates to non-reducing (but with boiling) SDS-PAGE (27). Under these conditions, abundant dimer formation was seen for WT NEMO, but not for the NEMO mutants Cys54/347Ala or 9SG (Fig. 3B). Given that Cys54 and Cys347 are intact in the 9SG mutant, its inability to form disulfide-linked dimers in the presence of hydrogen peroxide suggests that 9SG-NEMO has an altered conformation such that Cys54 and 347 are no longer suitably positioned within the noncovalent NEMO homodimer to form disulfide bonds

Overall, the results in this section show that the 9SG mutation alters the reactivity of cysteine residues in other regions of the protein, indicating that mutational disruption of the IVD alters the structure of the NEMO dimer in a way that propagates to distant parts of the protein.

### An inactivating mutation of IVD residues 145-153 perturbs ubiquitin-induced liquid-liquid phase separation of NEMO

Du et al. (18) and Goel et al. (19) have recently shown that ubiquitin can stimulate the LLPS transition of purified NEMO in vitro and that mutations that remove the C-terminal ubiquitin-binding domain of NEMO abolish this induced transition. Given our results above indicating that perturbation of the IVD interfered with the ability of NEMO to form higher order structures, we next determined whether IVD mutation affected LLPS of NEMO. Specifically, we compared LLPS formation of full-length and 9SG NEMO in the presence of M1-octa-Ubiquitin (M1-Ub_8_) (18). For the LLPS assay, we first purified full-length WT NEMO, 7XAla-NEMO, and 9SG-NEMO (7XAla background) expressed in *E. coli* and labeled the purified proteins with Alexa Fluor 488. The Alexa Fluor-labeled NEMO proteins were then mixed with increasing concentrations of M1-octa-Ubiquitin (M1-Ub_8_), and the reaction mixtures were monitored for liquid droplet formation by fluorescence microscopy. We found that both WT NEMO and 7XAla NEMO readily formed droplets in the presence of M1-Ub_8_, but few droplets were seen with 9SG-NEMO (Fig. 4A). The total raw fluorescence for both WT and 7XAla NEMO increased with increasing concentrations of M1-Ub_8_, whereas the total raw fluorescence of 9SG-NEMO did not increase under these conditions (Fig. 4B). In addition, at the same sub-saturating concentration of M1-Ub_8_, 7XAla showed approximately 2.5-fold more LLPS compared to WT NEMO. In contrast, 9SG-NEMO showed only approximately 10% of the total fluorescence of its positive control 7XAla-NEMO (Fig. 4C). The bacterially expressed proteins used in the LLPS assays is shown in Fig. 4D. Overall, these results demonstrate that the 9SG IVD mutation interferes with ubiquitin-induced LLPS of NEMO.

Since the 9SG mutation is not within the ubiquitin-binding region of NEMO, we sought to confirm that the effect of this IVD substitution on LLPS was not due to a failure of the protein to interact with ubiquitin. For these experiments, we incubated bacterially expressed GST alone or GST-diubiquitin (Ub_2_) with cell lysates from 293T NEMO KO cells overexpressing FLAG-tagged NEMO proteins and performed a pulldown assay as described previously (30). We used cell lysates expressing FLAG-tagged versions of WT, 7XAla, 9SG, or L153R NEMO, all of which have intact C-terminal Ub-binding domains. All of these NEMO proteins were pulled down by GST-Ub_2_, but not by GST alone (Fig. 4E). These results indicate that the inability of the 9SG NEMO mutant to undergo LLPS is not due to an impaired ability to bind to ubiquitin.

Taken together, the results in this section demonstrate that the core IVD sequence of NEMO is required for the LLPS that is induced by binding to ubiquitin in vitro.

### Mutation of IVD residues 145-153 perturbs the formation of induced puncta of NEMO in vivo

Previous studies have shown that induced activation of the NF-κB pathway in vivo can lead to the formation of higher order complexes containing NEMO, which can be observed as cytoplasmic puncta (31, 32). Furthermore, infection with mycoplasma has been shown to lead to constitutive activation of NF-κB in the chicken fibroblast cell line DF-1, due to chronic activation of upstream TLR6 (33). For our studies, we also chose to study induced puncta formation of NEMO proteins in DF-1 cells, in part because this cell line has an easily discernible “spread out” morphology.

We first confirmed that mycoplasma infection of DF-1 cells leads to constitutive activation of NF-κB by showing that infection of DF-1 cells with mycoplasma increased nuclear translocation of NF-κB p65 as compared to uninfected DF-1 cells (Fig. S6). We next used a surrogate for mycoplasma infection by treating cells with a synthetic lipoprotein mimetic, Pam2CSK4, which has been shown to cause NF-κB activation in other cell lines (34). As judged by nuclear translocation of NF-κB p65, Pam2CSK4 caused increased NF-κB activation following 4 h of treatment (Fig. 5A). We therefore used the 4 h time point for activation of NF-κB in our subsequent experiments.

**Figure 5.**
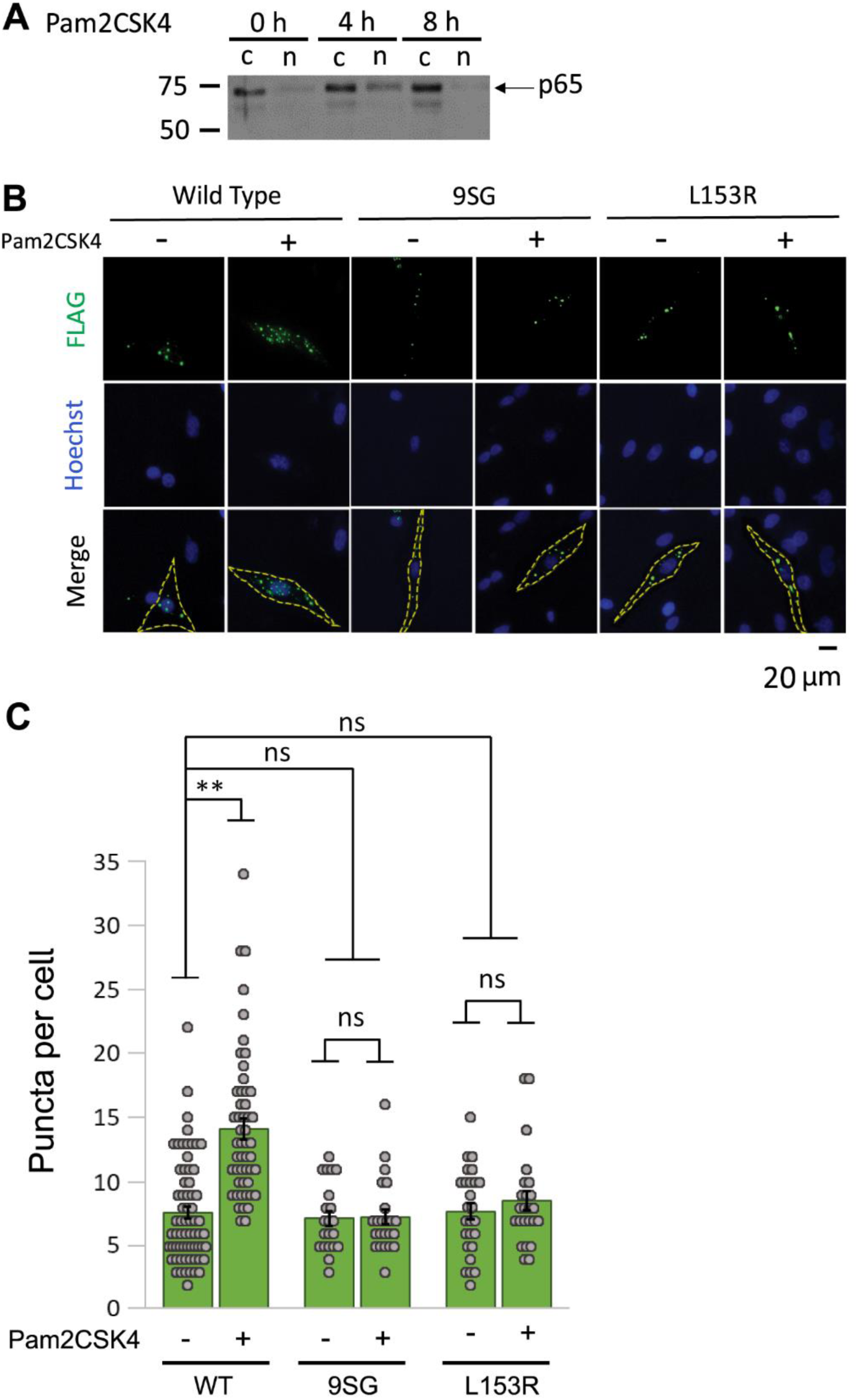
Inactivating mutation in the core region of the NEMO IVD affect puncta formation in vivo. (A) Anti-p65 Western blot of cytoplasmic (c) and nuclear (n) extracts from DF-1 cells treated for the indicated times with Pam2CSK at a final concentration of 10 µg/ml. (B) Representative fluorescent images of puncta formation in DF-1 cells transfected with the expression vectors for the indicated FLAG-tagged NEMO proteins. Cells were treated with 10 μg/ml Pam2CSK for 4 h (+) or were left untreated (-) prior to saponin extraction and fixation. (C) Quantitation of puncta per cell of transfected cells in (B). Shown are the means ± SEM; n = ≥ 20 cells per condition in at least two experiments. ns, not significant; **, p<0.001 by t-test assuming equal or unequal variance as appropriate. Dots are the number of puncta in individual cells.

To determine whether IVD perturbations affect NEMO puncta formation in DF-1 cells, we transfected DF-1 chicken fibroblasts with FLAG-tagged NEMO, 9SG NEMO or the human disease L153R mutant of NEMO. Two days later, transfected cells were treated with Pam2CSK4 for 4 h. We then performed anti-FLAG indirect immunofluorescence (IF) and counted the number of large cellular puncta (≥ 1 μm). Cells transfected with WT NEMO and treated with Pam2CSK4 had significantly higher numbers of puncta as compared to the corresponding untreated control cells, indicative of higher order complex formation, as consistent with previously published results (Figs. 5B, C) (31, 32). In contrast, cells transfected with the inactive 9SG and L153R NEMO mutants did not show increased numbers of puncta per cell following treatment with Pam2CSK4. Similar results were obtained by infection of cells with mycoplasma (Fig. S6). These results indicate that NEMO proteins with inactivating IVD mutations have an impaired ability to form higher order puncta in vivo following treatment with an activator of NF-κB and are consistent with the findings above that an inactivating IVD mutation reduces the ability of NEMO to undergo LLPS in vitro.

### Impact of the IVD on NEMO structure and stability

Previous work has suggested that the IVD acts to destabilize the coiled-coil structure of NEMO (15). To gain additional insight into the role of the IVD, and NEMO’s other domains, in the stability of the protein we generated a set of NEMO constructs with varying degrees of truncation N-terminally and/or C-terminally to the IVD (Fig. 6A). We tested the effect of these truncations on the thermostability of the protein using variable-temperature CD (Figs. 6B, 6D, S7). As described above, the T_m_ of full-length WT NEMO (1-419) was 40 ± 0.71 °C, which is consistent with previously published results (13). As expected, removal of the N-terminal disordered region from residues 1-43 (i.e., NEMO 44-419) had minimal effect on T_m_ (41 ± 0.23 °C), likely because this N-terminal region is largely unstructured (35).

**Figure 6.**
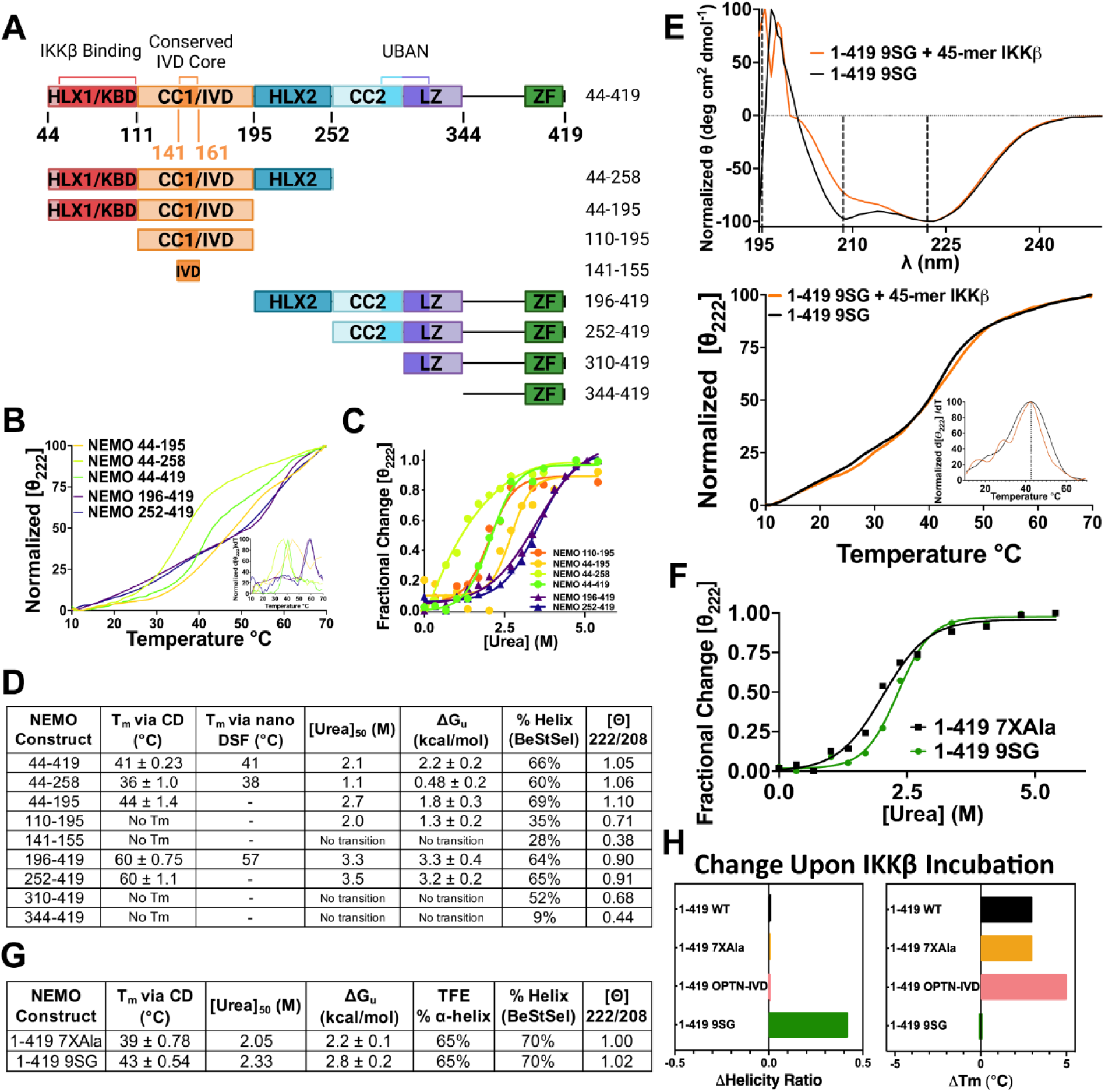
Impact of the IVD on NEMO structure. (A) Protein domain map showing the residues present in each truncated NEMO construct. (B) CD-monitored thermal denaturation of NEMO constructs measured by the loss in secondary structure monitored at 222 nm; inset showing the peak of the first derivative as the melting temperature (T_m_). (C) CD-monitored urea denaturation of NEMO constructs measuring the loss in secondary structure monitored at 222 nm (25 °C and 10 µM), fitted to an equation for two-state denaturation by nonlinear regression. (D) Tabulated values of truncated NEMO constructs for the following: T_m_ values determined via CD from (B) or via nanoDSF. The [urea]_50_ indicates the concentration of urea at which half of the protein is unfolded from (C). The ΔG_u_ values were calculated from linearized data in (C). The % helix was calculated from the CD spectra using the BeStSel server, and the ellipticity ratios were calculated from the signal at 222 nm to 208 nm. (E) CD experiments conducted with 1-419 9SG, ± equimolar equivalents of a 45-mer peptide of IKKβ, incubated for 1 h at room temperature. The top panel shows the CD spectra (25 °C; 10 µM) and the bottom panel shows the CD-monitored thermal denaturation with the inset showing T_m_ as in (B). (F) CD-monitored urea denaturation of NEMO measured as in (C). (G) Tabulated values for 1-419 7XAla NEMO and 1-419 9SG NEMO of the following: T_m_ determined by CD in (E); the [urea]_50_ in (F); the ΔG_u_ values calculated from linearized data in (F); the % alpha-helix calculated by comparing the 222 nm protein signal in TFE to that in buffer; and the % helix and ellipticity ratios calculated as in (D). (H) Bar plots comparing the Δhelicity ratio and ΔT_m_ for the indicated constructs ± IKKβ 45-mer peptide.

Of note, the constructs NEMO 196-419 and NEMO 252-419, which include only domains C-terminal to the IVD (and not the IVD itself), are considerably more stable (by 20 °C) than WT NEMO, having melting temperatures of T_m_ = 60 ± 1.1 °C for NEMO 252-419 and T_m_ = 60 ± 0.75 °C for NEMO 196-419. Thus, the presence of portions of NEMO N-terminal to residue 196 substantially destabilizes the protein in the presence of thermal denaturation, consistent with the notion that the IVD and/or the KBD exert a destabilizing effect on the HLX2 and the CC2/LZ domains which both comprise α-helical coiled-coils (29, 36).

Peptides comprising only the core conserved region of the IVD (NEMO 141-155), or even the full IVD (NEMO 110-195), do not show cooperative thermal unfolding events. Indeed, these small fragments are likely monomeric and disordered given that they lack the flanking domains that are mainly responsible for NEMO dimer stabilization (15). Dimerization has been observed by non-denaturing and native PAGE for constructs 44-419, 44-258, 44-195, 196-419, and 252-419 that include the flanking domains (data not shown). Inclusion of the adjacent KBD upstream of the IVD, in the construct NEMO 44-195 affords a fully cooperative thermal unfolding event, however, with T_m_ = 44 ± 1.4 °C, consistent with our prior work (13). A construct that includes the KBD and IVD plus the C-terminally adjacent HLX-2 domain (NEMO 44-258) also showed cooperative unfolding, having a T_m_ = 36 ± 1.0 °C, which is 4 °C lower than full-length NEMO and 8 °C lower than the KBD/IVD construct NEMO 44-195 (Figs. 6B, D). This result again shows that the KBD together with the IVD serve to destabilize “downstream” domains. Thermal denaturation results from nanoDSF corroborated these findings for select constructs (Fig. 6D).

Thermal denaturation of proteins is often irreversible, and thus a T_m_ value typically cannot be interpreted as a measure of thermodynamic stability as it is affected by kinetic factors arising from conditions such as protein concentration and temperature ramp rate. The observation of extensive precipitation upon heating in the variable temperature CD experiments described above established that thermal denaturation was indeed irreversible in these experiments. Therefore, we performed chemical denaturation experiments to determine the true thermodynamic stability of NEMO and its truncated variants (Figs. 6C, 6D, S8). In these experiments, the [urea]_50_ values represent the concentration of urea at which half of the protein is unfolded, and fitting the denaturation curve to the equation for a reversible, two-state denaturation model allows extrapolation to determine the Gibbs free energy of unfolding at zero urea (37, 38). The results of these experiments largely mirrored those from the thermal denaturation measurements. Specifically, the two constructs containing only those domains downstream of the IVD, i.e., NEMO 196-419 and NEMO 252-419, showed high free energies of unfolding of ΔG_u_ = 3.2-3.3 ± 0.2-0.4 kcal/mol, whereas NEMO 44-419, which additionally contains the KBD/IVD region, showed substantially reduced thermodynamic stability with ΔG_u_ = 2.2 ± 0.2 kcal/mol. Thus, the KBD and IVD collectively destabilize the NEMO dimer to the extent of ∼1.1 kcal/mol relative to NEMO constructs that contain only the HLX2 and CC2/LZ domains.

We additionally examined the CD spectra to assess the percent α-helix of the same constructs, and found a similar trend of the IVD decreasing the percent α-helix of the adjacent domains. When we calculated the percent α-helix from our CD spectra (Fig. 6D), we found that NEMO 44-419 and 44-195 have high α-helical content as we have previously shown (66% and 69%, respectively) (13). For the conserved core of the IVD (aa 141-155) alone, we measured a percent α-helix of 28%, which increases to 35% for the full IVD (110-195), and to 69% upon addition of the KBD and IVD domains (aa 44-195). The C terminus of NEMO (aa 344-419) has an extremely low percent α-helical content (9%) that increases as the LZ (310-419), CC2 (aa 252-419), and HLX-2 (aa 196-419) are added (52%, 65%, and 64%, respectively). When the IVD is added between the KBD and HLX-2 domains, a decrease in percent α-helix (to 60%) is observed, indicating a disruption in the secondary structure. These results are consistent with those from our T_m_ measurements and indicate that the IVD affects the conformation of adjacent domains.

To determine how much of this destabilization effect can be attributed to the IVD versus the KBD, we subjected 1-419 9SG (in the 7XAla NEMO background) to similar analyses. The CD spectrum of full-length 9SG (Fig. 6E) is similar to that of 7XAla (Fig. 2D), in agreement with our previous findings (13), showing that the loss of biological function seen for the 9SG mutant is not due to a major disruption of NEMO’s overall secondary structure. In the urea denaturation experiments, the free energy of unfolding for the 1-419 9SG mutant was ΔG_u_ = 2.8 ± 0.2 kcal/mol, compared to 2.2 ± 0.1 kcal/mol for the 7XAla control (Figs. 6F, G). These results show that disrupting the core region of the IVD by introducing the 9SG substitutions mitigated the stability difference caused by the KBD/IVD regions of NEMO, suggesting that this effect can be largely attributed to destabilizing interactions involving the IVD.

We also measured the effects of binding the IKKβ 45-mer peptide on the secondary structure and stability of different forms of NEMO. Addition of equimolar equivalents of the IKKβ peptide to WT NEMO, 7XAla-NEMO, or OPTN-IVD-NEMO in each case resulted in a substantial (3-5 °C) increase in thermal stability, as might be expected for addition of a high affinity ligand (39). No change in the coiled-coil content was observed upon IKKβ peptide binding to any of these three NEMO variants, consistent with the notion that the KBD region (aa 44-111) is preorganized as a coiled-coil even in the unbound state. In contrast, when 1-419 9SG was incubated with the IKKβ peptide, no T_m_ increase was observed, but there was a large increase in the ellipticity ratio from 1.02 to 1.44, indicating an increase in coiled-coil structure (Fig. 6H). This result suggests that the KBD region is disrupted in the unbound 9SG mutant, presumably because the disrupted IVD structure fails to correctly organize dimerization of the KBD, as has been shown to be necessary for proper coiled-coil formation (29, 40). Consequently, the binding energy that results from interaction with the IKKβ peptide is utilized to induce coiled-coil formation, with essentially none remaining to stabilize the protein. This conclusion is consistent with the differences between these NEMO constructs observed in prior small angle X-ray scattering analyses, which showed an overall lengthening of 9SG NEMO upon addition of the IKKβ peptide, in contrast with the compaction seen when WT NEMO bound the peptide (13).

## Discussion

The data presented herein further our understanding of the role of a conserved central region of NEMO (especially residues 145-153 of the IVD) that is required for full activity of NEMO in downstream activation of IKKβ and ultimately the activation of NF-κB. We show the extent of conservation of this region in NEMO proteins across a broad evolutionary swath, as well as sequence and functional conservation of the corresponding sequence from the NEMO-related protein optineurin (OPTN). Namely, we show that replacement of NEMO IVD sequences with the analogous OPTN sequences does not alter NEMO structure or function, whereas removal or replacement of aa 145-153 with a more flexible sequence (i.e., the Δ145-153 and 9SG mutants, respectively) rendered NEMO nonfunctional. The core IVD sequences are also required for LLPS in vitro and signal-induced puncta formation in vivo. Finally, we have used biochemical and biophysical methods to characterize effects that the IVD region has on NEMO structure, stability, conformational change, and higher order structure formation upon the binding of ligands, which lead to a model for how these attributes contribute to NEMO functionality. Overall, these results add to the mounting evidence that, during the course of NEMO’s scaffold signaling function, NEMO undergoes several conformational rearrangements upon binding various ligands, including IKKβ, IκBα, and poly-ubiquitin (12, 13, 15–17).

We previously suggested that the IVD was a mediator of conformational changes by showing that a compaction of NEMO occurs upon the binding of a 45-mer peptide derived from IKKβ and that mutation of residues 145-153 abolishes this effect (13). However, the lack of an experimental X-ray crystal structure or convincing structural model of the IVD left many questions unanswered about its architecture and function. Recent reports that NEMO engages in LLPS and that certain human disease mutants in NEMO that lack the ability to bind to linear ubiquitin are defective for LLPS in vitro, for the formation of higher order complexes in vivo, and for supporting induced NF-κB signaling (18, 19) add additional layers of complexity to NEMO’s mechanism of action.

How different conformational states are molecularly advantageous for NEMO function is not entirely known. One emerging model is that upon TNFα-induced activation of the NF-κB pathway, the NEMO-IKKβ complex is brought to the TNFR receptosome by binding to ubiquitin chains on the adaptor protein RIP. Further modification of NEMO with ubiquitin via LUBAC alters the conformational state of the NEMO complex, which is likely modulated, in part, by the IVD. This change in conformational state then provides a switch required for higher order IKK signalosome formation and LLPS of IKK signaling components, namely, NEMO, IKKβ, IκBα, TRAF, TAK/TAB, and LUBAC. Based on our biophysical data, we hypothesize that the IVD regulates the structure of NEMO by acting as a central hub to reorganize the flanking coiled-coil domains. This model is also consistent with observations by Ko et al. (15) who demonstrated that mutants lacking or containing mutations in the IVD exhibit altered solution dynamics, indicating perturbation in the ability of NEMO to oligomerize.

Our demonstration that disruption of the core sequences in the IVD (mutant 9SG) abolishes the ability of NEMO to undergo ubiquitin-induced LLPS coincides with the failure of this mutant to support activation of IKK in response to TNFα (11, 13). Moreover, we show that neither the 9SG mutant nor the L153R immunodeficiency disease mutant, which is also defective for IKKβ activation (11), form high molecular weight puncta in response to a TLR agonist. Thus, internal (e.g., IVD) mutations in NEMO also appear to disable NEMO signaling to IKK, by affecting LLPS and higher order signalosome formation, even though such mutants can still bind to Ub (Fig. 4E). Overall, we suggest that the 9SG and L153R mutations cripple NEMO function by disabling its ability to undergo a conformational change that is required for LLPS. Whether other NEMO human disease mutants—in the IVD or elsewhere—also affect LLPS is not clear. Our data are also consistent with puncta forming in stimulated cells via LLPS (18) in that an intact IVD core sequence is crucial for both LLPS of NEMO and for puncta formation in cells. Specifically, the 9SG mutation reduces LLPS, and the 9SG and L153R mutants do not show increased signal-induced puncta formation in cells.

Considering what we know about the conformational flux of NEMO upon ubiquitin binding and the solvent exposure of residues distal to the ubiquitin binding region (16), it is possible that LLPS of NEMO complexes is driven by the exposure of key residues that are buried residues in the resting state. A similar finding has been made for the scaffold HP1α that undergoes LLPS with heterochromatin. HP1α also has a low complexity hinge region that is important for LLPS, which is driven by ion concentrations that cause conformational changes in HP1α. It has been proposed that this reorganization of HP1α leads to solvent exposure of hydrophobic/charged residues and enhances self-oligomerization, which then promotes LLPS (41). It is presently unclear whether the deficiency in IκBα phosphorylation that is seen with the 9SG and L153R mutants (11, 13) is a result of these defects in LLPS or instead represents a different consequence of the disruption of IVD structure and function.

Our biophysical results also shed new light on the molecular details of how the IVD performs its critical functions. The results of the thermal and chemical denaturation experiments establish that the IVD mediates an intimate allosteric communication between the flanking KBD and the HLX2 and CC2/LZ domains. Evidence that the IVD is allosterically coupled to the HLX2 and CC2/LZ domains is provided by the observation that the presence of the KBD and IVD substantially destabilizes the protein compared to constructs in which these domains are deleted, namely NEMO 195-419 and NEMO 252-419, and this destabilization is largely relieved if the structure of the IVD is modified by replacement of its highly conserved core region with a flexible linker, in the 9SG mutant. These findings could be taken as evidence that the homodimer interface in the IVD region is intrinsically destabilizing. However, our results indicate that this is not the case. Inclusion of the IVD induces the otherwise largely disordered KBD region to adopt an ordered structure with substantial thermal stability, as shown by the thermal and chemical denaturation results for the KBD-IVD construct NEMO 44-195 as compared to the behavior of constructs containing only the KBD (7, 29, 35). This result shows that the IVD region can stabilize a homodimeric structure, and that NEMO’s KBD and IVD form a single structural unit that undergoes a cooperative unfolding transition, at least in the context of the truncated construct NEMO 44-195. The IVD is only seen to be destabilizing when the downstream HLX2 and CC2/LZ domains are also present, as, for example, in full-length NEMO. Our findings imply that the IVD has an organized three-dimensional structure with an interface between the two chains of the NEMO homodimer that is not intrinsically destabilizing, but in the context of full-length NEMO the IVD becomes enthalpically strained so as to destabilize the protein. This strain can be explained by conflicting geometric or conformational demands placed on the IVD by the upstream KBD and the downstream HLX2/LZ domains. We propose that substituting the core region of the IVD with a flexible 9SG linker relieves this strain by disrupting the structural integrity of the IVD such that it can now simultaneously conform to the geometric preferences imposed by the KBD on one side and by the HLX2/CC2/LZ domains on the other. Thus, the IVD serves to balance the conflicting conformational requirements of the flanking regions of NEMO, placing the two ends of the protein in allosteric communication with each other.

One explanation for how this strain is imparted to the IVD is that the IVD, like its neighboring domains, normally exists as an α-helical coiled coil, but is capable of adopting two different intra-dimer interaction angles between the helices. If one twist angle between the IVD helices were most compatible with the KBD dimer structure, but a different twist angle fit better with the constraints imposed by the downstream HLX2/CC2/LZ domains, then the intra-chain interactions within the IVD could be stabilizing in a KBD-IVD construct, as we observed experimentally, but would be destabilizing in the full-length protein as the IVD cannot satisfy the conflicting geometric demands of the HLX2/CC2/LZ domains. As an alternative to this “IVD helical twist” model, it could be that, in the KBD/IVD construct NEMO 44-195, only the upstream parts of the IVD form a stabilizing coiled-coil interaction whereas the downstream IVD segments are disordered or do not contact each other. In this scenario, when the HLX2/CC2/LZ domains are present they force an association between the downstream regions of the IVD that enforce energetically unfavorable contacts. Other models to account for how strain in the IVD mediates allosteric communication between the KBD and the HLX2/LZ domains might also be imagined.

One key question is how the accumulation of strain energy in the IVD is utilized in NEMO’s function. The use of binding energy derived from favorable interactions in opposition to destabilizing interactions elsewhere in the same protein is known to be a means by which proteins achieve and regulate their functions (42, 43). We previously showed that binding of an IKKβ-derived peptide to the KBD of NEMO induces a conformational change in NEMO (13). In light of the mechanism of IVD function proposed herein, this previous finding can be interpreted as IKKβ peptide binding acting to lock the KBD into the rigid geometry that is observed experimentally (35), and thereby modulating the geometric constraints on the nearby IVD. It is plausible that altering the geometry of these portions of the highly strained IVD causes a change in IVD conformation, in this case resulting in a “kink” in the protein, as seen in the IVD region by SAXS (13). The key role that IVD strain plays in this conformational change is shown by the observation that introduction of the flexible 9SG mutation abolishes this conformational effect of IKKβ peptide binding (13). In cells, the association of NEMO with IKKβ is likely constitutive, based on the high affinity of this interaction (23), and therefore, the conformational change in NEMO that results from IKKβ binding is unlikely to reflect a process that is part of NEMO’s signaling mechanism. However, it provides direct evidence for an intimate allosteric connection between the KBD and IVD, and for the importance of the IVD in determining the overall conformation of the NEMO dimer. Based on the allosteric communication that appears to exist between the KBD and the HLX2/LZ domains, we speculate that the conformational effects on NEMO seen upon binding of polyubiquitin to the UBAN site in the LZ domain (12, 17) are also modulated by the IVD.

Our finding that the critical core region of the NEMO IVD can be substituted by the corresponding sequence from OPTN, with full retention of structure as measured by CD and disulfide-mediated dimerization assays and with full function as measured in cellular signaling assays, suggests that the IVD-like region of OPTN is both structurally and functionally analogous to the NEMO IVD. NEMO and OPTN have considerable overall aa sequence similarity and a similar domain organization, and in some cases OPTN can compete with NEMO for binding to ubiquitin (44, 45), but their biological functions are largely distinct. That is, OPTN is a membrane-bound protein that has primarily been ascribed roles in autophagy, as a cargo receptor that transports ubiquitinated targets to lysosomes for degradation. Like NEMO, mutations in OPTN have been found in several human diseases, but the OPTN-associated diseases are neurological diseases including ALS and glaucoma (46). Of note, two ALS mutations fall within the core region of the IVD-like region of OPTN that we used for our substitution experiments (46). Whether the IVD-like core sequence in OPTN also promotes LLPS is not known. It is of note, however, that OPTN may be part of pathogenic aggregates that form in ALS neurons (47).

One curious finding is that although single-residue human disease mutants have been found along the length of the NEMO protein (5), the 7XAla-NEMO mutant–which has seven Cys residues converted to Ala, spanning the region from aa 11 to 347–appears to be fully functional and structurally intact (28). The 7XAla-NEMO protein is useful for biochemical and biophysical analyses because, unlike WT NEMO, bacterially expressed 7XAla NEMO does not form disulfide-bonded aggregates (28). We also found that 7XAla-NEMO forms a greater number of ubiquitin-induced LLPS droplets in vitro (Fig. 4B). One possibility is that 7XAla-NEMO (in the absence of disulfide bonds) can more readily transition to conformational states that promote LLPS upon binding to ubiquitin. Our comparative sequence analysis showed that the seven Cys residues are generally conserved in mammals (Fig. S3), however, they are not present in NEMO from many non-mammalian species (data not shown), which is consistent with our demonstration that these Cys residues are not essential for NEMO activity (28, 29).

A limitation of our studies is that in our in vitro experiments we have examined the structure of NEMO in isolation or when bound to single ligands (e.g., ubiquitin or an IKKβ peptide). Clearly, in vivo NEMO, as a scaffold, binds to multiple protein substrates in vivo, including IKKβ, LUBAC, different poly-ubiquitins, TRAF proteins, IκBα, and perhaps others (20, 31, 48–50). Furthermore, in cells, NEMO is subject to other context-dependent modifications, such as phosphorylation (51, 52). All of these interactions and modifications are likely to affect the conformational states of NEMO. Nevertheless, because the conformational plasticity of NEMO is at the heart of dynamic events such as LLPS and signaling, understanding how different NEMO sequences contribute to conformational plasticity provides important fundamental scientific knowledge.

As we have recently reviewed (1), NEMO is one of several scaffold proteins that have been demonstrated to undergo context-dependent conformational changes that are important for directing and optimizing signaling events. Similarly, LLPS is important for many cellular and biomolecular events (53). Moreover, human disease mutations that affect dynamic structural changes in some other scaffolds have been documented (1), as we suggest herein for mutations within the IVD of NEMO. Our results are consistent with a picture in which the IVD acts as a central hub that regulates the orientation of the flanking α-helical, coiled-coil domains that interact with key binding partners, and places these N- and C-terminal regions of the protein in allosteric communication with one another. Mutations in the IVD can destroy the structural integrity of this hub, and thereby impair the overall function of NEMO. Therefore, understanding how these conformational changes occur and how they are affected in disease could lead to the development of new types of therapeutics. Such strategies may be analogous to what has been done for the therapeutic stabilization of common mutations in the cystic fibrosis transmembrane conductance regulator (54).

## Experimental procedures

### Evolutionary rate, hydropathy and similarity network analyses

A BLAST database search was conducted for nonredundant protein sequences in the NCBI database using the human NEMO sequence as the query, yielding 500 sequences from vertebrate species with sequence identities ranging from 64-100%. The sequences were aligned using Clustal Omega (EMBL-EBI) (55) and formatted for input to the program Rate4Site (22) version 3.0.0, which was run using default settings to create a phylogeny tree, and then to calculate the evolutionary rate of each specific residue, with the output plotted and visualized using a four-period moving average.

The hydropathy of each individual residue was scored according to the Kyte & Doolittle hydropathy scale, plotted, and then subjected to a moving average for visualization. The evolutionary rates of ordered regions of NEMO with known structure were averaged, as were the evolutionary rates of uncharacterized regions of interest, which were divided by the ordered region average to obtain the indicated ratios.

For multiple sequence alignment (Fig. 1D), a BLAST database search using human NEMO as a reference sequence was conducted using Clustal Omega (EMBL-EBI). The alignment visualized and colored according to residue conservation using MView (56).

For the SSN analysis (Fig. S2), an initial sequence BLAST search was conducted using human NEMO with a UniProt query e-value threshold of 1 x 10^-5^, with filtering to remove fragments and non-eukaryotic sequences using the EFI-EST tool. A final alignment score cut-off of 61% was used to obtain the set of sequences to conduct the SSN analysis with an e-value threshold of 1 x 10^-60^, and 674 sequences and 2708 nodes were imported into the program Cytoscape (57) for manipulation and visualization.

### Plasmids

The pcDNA-FLAG expression vector has been described previously (11). cDNAs encoding WT and mutant IVDs were synthesized (GenScript or TWIST Bioscience) or generated via PCR (Supplemental Table 1). The pcDNA-7XAla and 9SG variants have been described previously (13). These cDNAs and PCR-generated IVD mutants were subcloned into the appropriate vector using the cloning strategy outlined in Supplemental Table 1. All plasmids were purified with a Midiprep kit (ZymoPURE, #D413) or a QIAprep spin miniprep kit (QIAGEN, #27104) and verified by DNA sequencing prior to use. Primers used for PCR amplification are included in Supplemental Table 2. SUMO-NEMO-IVD clones were prepared in the pE-SUMOstar vector (Life Sensors PE-1106-0020) using the cloning strategies outlined in Supplemental Table 1.

### Cell culture, transfection, and cell treatments

Human HEK 293T, HEK 293T NEMO KO, and DF-1 chicken fibroblast cells were grown in Dulbecco’s modified Eagle’s Medium (Invitrogen) supplemented with 10% fetal bovine serum (Biologos), 50 units/ml penicillin, and 50 mg/ml streptomycin at 37°C, 5% CO_2_ in a humidified chamber as described previously (11, 58). Transfection of cells with expression plasmids was performed using polyethylenimine (PEI) (Polysciences, Inc.) as described previously (29, 58) or with Effectene transfection reagent (Qiagen, 301425) according to the manufacturer’s instructions, as described previously (11). Briefly, for PEI transfection, cells were incubated with plasmid DNA and PEI at a DNA:PEI ratio of 1:6. Media was changed 24 h post-transfection, and whole-cell lysates were prepared 24 h later in AT Lysis Buffer (20 mM HEPES, pH 7.9, 150 mM NaCl, 1mM EDTA, 1 mM EGTA, 20% w/v glycerol, 1% v/v Triton X-100, 20 mM NaF, 1 mM Na_4_P_2_O_7_·10H_2_O, 1 mM dithiothreitol (DTT), 0.2% Protease Inhibitor Cocktail [Sigma-Aldrich, P8340]).

For NF-κB translocation and NEMO puncta formation analyses, DF-1 chicken fibroblasts were stimulated with the synthetic TLR2/6 antagonist Pam2CSK4 (InvivoGen tlrl-pms) at a final concentration of 10 µg/ml for the indicated times prior to lysis or fixation. Infection of DF-1 cells with mycoplasma was verified by the ATCC mycoplasma detection kit (ATCC 30-1012K) (Fig. S6). For analysis of phosphorylation of IκBα, 293T NEMO KO cells were stimulated with TNFα at a final concentration of 20 ng/ml for 10 min prior to lysis. To assess the formation of oxidation-induced NEMO dimers, cells were treated with 50 µM H_2_O_2_ for 10 min prior to lysis.

### Western blotting

Whole-cell extracts from 293T cells were prepared as described previously (11, 58) in AT lysis buffer (20 mM HEPES pH 7.9, 150 mM NaCl, 1mM EDTA, 1 mM EGTA, 20% w/v glycerol, 1% v/v Triton X-100, 0.2% Protease Inhibitor Cocktail [Sigma-Aldrich P8340]). Samples analyzed under standard reducing conditions were heated at 95°C for 10 min in a loading buffer containing 5% β-mercaptoethanol prior to SDS-PAGE. Samples analyzed under non-reducing conditions did not contain β-mercaptoethanol and were either heated (Fig. 3B) or not heated (Fig. 3A) prior to SDS-PAGE. In all cases, extracts were separated on 10% SDS-polyacrylamide gels, and then electrophoresed onto a nitrocellulose membrane in transfer buffer (20 mM Tris, 150 mM glycine, 20% methanol) overnight at 170 mA, 4 °C. Membranes were blocked in TMT (10 mM Tris-HCl pH 7.4, 150 mM NaCl, 0.1% v/v Tween 20, 5% w/v Carnation nonfat dry milk) for 1 h at room temperature, and incubated in the appropriate primary antibody overnight at 4 °C. Primary antisera were as follows: anti-NEMO (1:2000; #2689, Cell Signaling Technology); anti-phospho-IκBα (1:1000; #9246, Cell Signaling Technology); anti-p65 (1:2000, #1226 gift of Nancy Rice); and anti-FLAG (1:1000, #2368S, Cell Signaling Technology). Membranes were washed four times with TBST (10 mM Tris-HCl pH 7.4, 150 mM NaCl, 0.1% v/v Tween 20). Membranes were then incubated with the appropriate secondary HRP-linked antibody as follows: anti rabbit-HRP (1:3000, Cell Signaling Technology 7074) for NEMO, FLAG, tubulin, p65 or anti-mouse-HRP (1:3000, Cell Signaling Technology 7076) for anti-phospho-IκBα. Membranes were then washed three times with TBST, twice with TBS (10 mM Tris-HCl pH 7.4, 150 mM NaCl), and incubated for 5 min at room temperature with chemiluminesent HRP substrate (SuperSignal West Dura Extended Duration Substrate, Thermo Fisher).

Immunoreactive bands were detected with a Sapphire Bimolecular Imager. For determination of NEMO activity, we measured the phosphorylation of IκBα in response to TNFα stimulation for all constructs in triplicate separately performed transfections. An F-test was first performed to determine the equality of variance, and then either a Student’s t-test, or a Welch’s t-test was performed to determine significance, based on the equality of variance.

### Indirect immunofluorescence

Transfection and indirect immunofluorescence of DF-1 cells was performed essentially as described previously (30, 58). Briefly, transfected DF-1 cells were seeded onto glass coverslips at an approximate density of 10^5^ cells/coverslip and grown overnight. Cells were then washed twice with ice-cold PBS and then twice with saponin extraction buffer (80 mM Pipes, pH 6.8, 1 mM MgCl_2_, 1 mM EGTA, 0.1% saponin) for 10 min to select for higher order NEMO complexes (18, 32). Cells were then fixed with 4% paraformaldehyde for 15 min at room temperature, blocked with blocking buffer (3% normal goat serum, 0.2% Triton-X-100 in PBS) for 1 h, and probed with an anti-FLAG primary antibody (1:100, #2368S, Cell Signaling Technology) diluted in blocking buffer for 2 h. Coverslips were washed three times with PBS for 5 min each, and probed with an Alexa Fluor 488-conjugated goat-anti-rabbit secondary antibody diluted in blocking buffer (1:100, #A11008, Invitrogen) for 1 h at 37°C. Coverslips were then washed three times with PBS for 5 min each, counterstained with Hoechst (1:10,000, B2883-25MG, Sigma-Aldrich) for 1 min before mounting onto glass slides with SlowFade Gold antifade reagent (S36937, Invitrogen). Cells were examined under an Olympus IX70 inverted fluorescence microscope using an Olympus DPlanApo 40X UV objective. Images were analyzed in ImageJ utilizing Otsu thresholding and counting the number of puncta with a diameter of at least 1 µm. An F-test was first performed to determine the equality of variance, and then either a Student’s t-test, or a Welch’s t-test was performed to determine significance, based on the equality of variance.

### Protein expression in bacteria and purification

Bacterial expression plasmids for full-length (aa 1-419) WT NEMO, 7XAla NEMO and 9SG NEMO coding sequences were transformed into Rosetta 2(DE3) pLysS competent cells (Novagen) as previously described (28). For NEMO SUMO 44-195, NEMO SUMO 110-195, NEMO 44-419, NEMO 44-258, NEMO 196-419, and NEMO 1-419 OPTN-IVD, plasmids were transformed into T7 Express DE3 *E. coli* competent cells (New England Biolabs) as previously described (13, 28). NEMO SUMO 252-419, NEMO SUMO 310-419, and NEMO SUMO 344-419 were transformed into BL21(DE3) *E. coli* competent cells (New England Biolabs). The SUMO tags were cleaved from SUMO-tagged proteins prior to use in experiments.

All *E. coli* cells containing vectors for bacterial expression of NEMO proteins were plated on Luria broth (LB) agar plates containing the appropriate antibiotic and were incubated overnight at 37 °C. A single colony was picked and used to inoculate 50 ml of LB containing the appropriate antibiotic(s) and incubated overnight at 37 °C with shaking at 200 rpm. Five ml of the overnight culture was used to inoculate 1 liter of LB containing the appropriate antibiotic(s) and incubated at 37 °C with shaking at 250 rpm until the culture reached an OD_600_ of ∼0.6. Isopropyl β-D-1-thiogalactopyranoside (IPTG) (GoldBio) was then added to a final concentration of 1 mM, and cells were grown for another 4 h. Cells were then isolated by centrifugation at 6000 x g for 15 min at 4 °C, the supernatant was discarded, and the pelleted cells were stored at -80 °C.

For purification, pelleted cells from 4 L of LB were thawed for 30 min at room temperature and solubilized in 60 ml of lysis buffer containing 20 mM sodium phosphate (Sigma-Aldrich) pH 7.5, 500 mM NaCl (RPI Research Products International), 40 mM imidazole (RPI Research Products International), two EDTA-free Pierce protease inhibitor tablets (Thermo Fisher), 600 μl of 10 mg/ml DNAse I (GoldBio) with all purification buffers for full-length WT NEMO also containing 5 mM Tris(2-carboxyethyl)phosphine (TCEP) (GoldBio) to minimize the formation of disulfide0bonded aggregates. Resuspended cells were lysed using a M110P microfluidizer (Microfluidics) at 18,000 psi and 8 M urea was added and incubated with stirring for 1 h at room temperature to dissociate contaminating proteins and prevent them from copurifying with NEMO. The cell debris was pelleted by ultracentrifugation at 35,000 rpm for 30 min at 4 °C, and then the supernatant was sonicated to shear remaining DNA fragments. The remaining purification steps were carried out at 4 °C using an ÄKTA protein purification system (GE Healthcare/Cytiva) for column chromatography steps. Specifically, the clarified lysate was added to a 5 ml HisTrap High Performance column (GE Healthcare/Cytiva), washed with 5 column volumes of lysis buffer supplanted with 6 M urea, with the bound protein refolded on-column using a urea gradient from 6 to 0 M urea over 20 column volumes. The protein was eluted using an imidazole gradient made with lysis buffer that was supplanted with 500 mM imidazole. The eluted protein was concentrated using Amicon Ultra-15 Centrifugal Filter Units (Millipore-Sigma) centrifuged at 4,000 x g to no more than 3 ml, and then purified using size exclusion chromatography (SEC) to remove contaminating proteins, NEMO dimers and other higher order oligomers that were present during purification of constructs containing significant portions of the coiled-coiled dimerization domains. The SEC was performed using a HiPrep 26/60 Sephacryl S-300 HR column (GE Healthcare/Cytiva) in storage buffer containing 20 mM sodium phosphate pH 7.5, 500 mM NaCl, with a flow rate of 0.6 ml/min, and collected in 3 ml fractions, Fractions containing monomeric NEMO were pooled, concentrated as above to ∼1-5 mg/ml, flash frozen in liquid nitrogen, and stored at -80 °C. For NEMO constructs containing an N-terminal SUMO fusion protein, the SEC step was omitted. Instead, the SUMO tag was removed, by first concentrating the eluate from the HisTrap column as above to 1 ml, diluting 5X with storage buffer containing 1 mM DTT (GoldBio), and incubating with either 2.5 µl (∼25 units) of SUMOstar Protease 1 (LifeSensors) for constructs in SUMOstar vectors or with SUMO Protease (Thermo Fisher) for constructs in non-SUMOstar vector for 4 h at 25 °C followed by incubation at 4 °C overnight. The reaction mixtures were then loaded onto a 5 ml HisTrap High Performance column, and the flow-through fractions containing the untagged NEMO proteins were collected. concentrated, and flash frozen as described above. The purifications of NEMO 252-419, NEMO 310-419, and NEMO 344-419 were conducted without the denaturation step, as contaminating proteins copurifying with other NEMO constructs were not visible during the purification of these constructs.

Protein concentrations for the constructs 1-419 WT, 1-419 7XAla, 1-419 9SG, 1-419 OPTN-IVD, and 44-419 were calculated from absorbance at 280 using the calculated extinction coefficient of each construct, calculated with the ProtParam tool on the Expasy web server (59) and measured using a NanoDrop 2000 (Thermo Fisher). For NEMO 110-195, the concentration was calculated by means of the Lowry protein assay using Folin & Ciocalteu Reagent (Thermo Fisher). For all other proteins, the concentration was calculated by the Bradford assay using a Quick Start Bradford Protein Assay Kit 2 (Bio-Rad).

### SDS-polyacrylamide gel electrophoresis of bacterially expressed proteins

Samples analyzed by SDS-PAGE were separated under standard reducing conditions and were boiled at 95°C in a loading buffer containing a final concentration of 5% β-mercaptoethanol. Proteins were then separated on 10-20% SDS-polyacrylamide gels and were visualized by staining with Coomassie blue (Bio-Rad).

### Peptide synthesis and purification

#### Peptide synthesis

The peptide fragment corresponding to NEMO residues 141-155 was synthesized using by Fmoc solid phase fast-flow peptide synthesis (60) with an N-terminal acetyl GC group. The reactor was filled with 190 mg of H-Rink Amide-ChemMatrix resin (Biotage) and swelled with dichloromethane (DCM) (Thermo Fisher, certified ACS) to help remove air bubbles, sealed, connected to the HPLC pump set at 8.3 ml/min, and rinsed with N,N-Dimethylformamide (DMF) (Thermo Fisher certified ACS) for several min while upside down with agitation until there were no visible air bubbles. The flow was then halted and the reactor placed in a 70 °C water bath and allowed to equilibrate to 70 °C for 5 min, followed by a two-min flush with DMF. A 0.4 M solution of 2-(7-aza-1Hbenzotriazole-1-yl)-1,1,3,3-tetramethyluronium hexafluorophosphate (HATU) (Chem-Impex International) was prepared in DMF and 2.5 ml was used to dissolve 1 mmol of each Nα Fmoc-protected aa (Chem-Impex International). Immediately prior to use, the aa solutions were activated with 1.56 eq of N,N-diisopropylethylamine (DIPEA) (Sigma-Aldrich, Reagent Plus, ≥99%), excepting cysteine where only 1.1 eq of DIPEA was used. Each activated aa was injected using a syringe pump at 8.3 ml/min for 30 sec, for the coupling step, followed by a 40 sec wash with DMF, a 20 sec deprotection step with 20% piperidine (Sigma-Aldrich, Reagent Plus, 99%) in DMF to remove the Fmoc groups, and lastly a 1-min wash step with DMF. This process was repeated for each aa addition, followed by injection of 10 eq of acetic anhydride plus 10 eq of DIPEA in DMF, injected using a syringe pump at 8.3 ml/min for 36 s to acylate the N-terminus, and then a final 2-min wash with DMF. The resin, containing the acylated peptide, was dried inside the reactor by pumping air through it. Then, the resin was removed, placed in a fritted syringe (Torviq), rinsed five times with DCM and two times with methanol, and dried under vacuum for 30 min.

#### Peptide cleavage from resin

The dried resin was transferred to a 15 ml conical tube, and a 10 ml cocktail of 2.5% water, 2.5% triisopropylsilane (TIS) (Sigma-Aldrich), 2.5% ethane-1,2-dithiol (EDT) (TCI America), and 9.25% trifluoroacetic acid (TFA) (Sigma-Aldrich) was added, followed by incubation for 3 h at room temperature with agitation. The solution was added to a fritted syringe and rinsed with 15 ml TFA into a 50 ml conical tube. The flow-through containing the peptide was dried under a N_2_ (g) stream until approximately one ml remained, at which point 15 ml of diethyl ether (Sigma-Aldrich, anhydrous, ACS grade) previously chilled to 4 °C was added, following by vigorous shaking, and the tube was cooled in N_2_ (l) for 2 min. The tube was centrifuged at 6,000 x g at 4 °C for 15 min, the supernatant was discarded, and the tube was left to dry at room temperature for 4 h.

#### Peptide purification

The successful synthesis and cleavage of the peptide was confirmed on a test sample using liquid chromatography/mass spectrometry (LCMS). Liquid chromatography separation was performed with a gradient between mobile phase A of water with 0.1% formic acid (Thermo Fisher, 99.0+%, Optima LC/MS Grade) and mobile phase B of acetonitrile (Thermo Fisher, HPLC grade) with 0.1% formic acid, using a 30 mm by 2.1 mm C18 column with 3 μm sized particles (Waters) using a Q-TOF Premier LCMS system (Waters). The dried peptide was then dissolved in 1.5 ml of 50:50 water:acetonitrile and purified using a 25 g C8 column (Interchim) on a puriFlash XS520Plus system (Interchim), with a mobile phase A of water with 0.1% TFA and a mobile phase B of acetonitrile with 0.1% TFA, and collected in fractions. The fractions were checked by LCMS to identify those containing pure peptide, and these fractions were pooled, dried using a centrifugal Genevac EZ-2 Personal Evaporator, and stored at -20 °C. Peptide concentrations were determined by the Lowry protein assay using Folin & Ciocalteu Reagent (Thermo Fisher).

### Protein labeling and in vitro liquid-liquid phase separation assay

Bacterially expressed and purified NEMO proteins were labeled with the Alexa Fluor 488 Microscale Protein Labeling Kit (Thermo Fisher, A30006) according to the manufacturer’s instructions, using a molar ratio of 10:1 for Alexa Fluor® 488:NEMO. The final degree of labeling of NEMO proteins was calculated to be approximately 2 molecules of Alexa Fluor® 488 dye per molecule of NEMO.

To measure phase separation of NEMO proteins, 6 µM of labeled NEMO protein was incubated with concentrations from 0-2.25 µM of M1-Ub_8_ (#BML-UW0805-0100, Enzo Life Sciences) in a buffer containing 20 mM Tris-HCl, pH 7.5, 150 mM NaCl, 5 mM ATP, 2 mM DTT, and 0.2 mg/ml BSA. Mixtures ((total volume 20 µl) were incubated at 37°C for 30 min in 384-well plates (that were previously coated for 1 h with 30 mg/ml BSA at 37 °C) (#3544, Corning). Imaging was then performed on a Nikon C2+ confocal microscope equipped with a thermal chamber set to 37°C. Fluorescence intensities were measured by quantifying the total fluorescence using ImageJ.

### Glutathione-S-transferase (GST) pull-down assays

GST and GST-Ub_2_ domain fusion proteins were expressed in BL21 *E. coli* cells and were purified from extracts using glutathione agarose (#16100, Thermo Fisher) as described previously (61). The GST proteins on glutathione beads were incubated with extracts from 293T NEMO KO cells expressing the indicated FLAG-tagged NEMO proteins (0.5% of each cell lysate was used as an input control). The beads were then washed four times with cold PBS, boiled at 95°C for 10 min in SDS sample buffer containing 5% β-mercaptoethanol, and electrophoresed on a standard denaturing 10% SDS-polyacrylamide gel. Pulled-down proteins were visualized by anti-FLAG Western blotting as described above (30, 58) (using anti-FLAG antiserum at 1:1000, #2368S, Cell Signaling Technology). To assess the amounts and sizes of the purified GST proteins used in the pulldowns, 1% of GST samples were electrophoresed on a 10% SDS-polyacrylamide gel, which was then stained with Coomassie blue (Bio-Rad).

### Circular dichroism spectroscopy, and α-helical content and coiled-coil content determination

The different purified NEMO proteins were diluted to 10 uM with 20 mM sodium phosphate buffer containing 500 mM NaCl at pH 7.4. CD spectra were obtained using an Applied Photophysics Chirascan CD spectrometer with a Starna Spectrosil quartz cuvette, 1 mm path length, under constant nitrogen flush. Buffer spectra were recorded from 190 to 260 nm with a 0.5 nm step at 0.5 s/step and at a temperature of 25 °C. The raw signal, measured in millidegrees, was converted to mean residue ellipticity (deg cm^2^ dmol^-1^), dividing by the number of residues (62). The ellipticity ratio used to evaluate the coiled-coil content of each construct was calculated from the ratio of the mean residue ellipticity measured at 222 nm to that at 208 nm. To determine alpha helical content, each construct was concentrated to 335 μM, diluted to 10 μM with 97% with 2,2,2-trifluoroethanol (TFE), and the CD spectrum measured at 222 nm after incubation for 2 h at room temperature. This signal was compared to that of the construct similarly diluted and incubated, but in CD buffer rather than TFE, using Equation **1**.

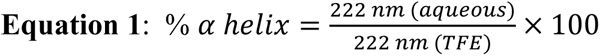

### Thermal denaturation

Melting temperatures (T_m_) were determined by monitoring the thermal denaturation of the proteins by CD. The raw signal at 222 nm was measured in millidegrees as the temperature was increased from 10 to 70 °C at a rate of 1 °C/min at a step of 0.2 °C, with a tolerance of 0.15 °C, a collection time of 0.5 sec per time point, and was converted to mean residue molar ellipticity. The T_m_ values were taken as the maximum of the first derivative of the thermal melting curves as a function of temperature. The reported T_m_ values represent the average of at least 3 independent measurements, ± the standard deviation.

### Urea denaturation

The purified NEMO proteins were concentrated to 100 µM and then diluted 10-fold with CD buffer supplemented with varying concentrations of urea. The samples were incubated at RT for 60 min to allow unfolding to occur. Incubation times of several hours were first tested to ensure that 60 min was sufficient time for equilibrium to occur. CD spectra of the samples were collected at 25 °C from 190 to 260 nm in 0.5 nm steps, measured for 0.5 sec/step, using buffer containing the same urea concentration but without protein for background subtraction. The denaturation data were plotted after conversion from units of mdeg to MRE as above, and fitted to the equation for two-state denaturation by nonlinear regression as previously described (29). The reported values for ΔG_u_ represent the average of the results from at least two independent experiments.

To test whether the urea-induced denaturation was reversible, for each construct a 100 µM sample in CD buffer was diluted 1:1 with CD buffer containing 8 M urea and incubated for 60 min at RT. The samples were then diluted to 10 µM in CD buffer without urea followed by incubation for a further 60 min to allow refolding to occur. In each case, a control was conducted using CD buffer without urea for the denaturation step. CD spectra were measured as described above and compared to ensure the NEMO constructs were able to fully refold.

### NanoDSF

Nano differential scanning fluorimetry data were collected using a NanoTemper Prometheus NT.48 instrument. NT.48 series nanoDSF grade standard capillaries (Nanotemper) were loaded with ∼25 μl of protein at 1-5 mg/ml in 20 mM sodium phosphate buffer, pH 7.4, 500 mM NaCl. The tryptophan and tyrosine residues were excited at 285 nm using 100% excitation power, and the resulting fluorescence was measured at 330 and 350 nm while the temperature of the sample was ramped from 20 to 70 °C at a rate of 1 °C/min. Using the included PR ThermControl software, the fluorescence data were plotted as a function of temperature and fitted to an arbitrary polynomial function. To determine the melting temperature, the first derivative of the ratio of the 350 to 330 nm curves was taken.

## Supporting information

This article contains supporting information.

## Acknowledgments

This work was supported by NIH grant GM117350 (to A.W.). J.W. and J.J.J. were supported by the Boston University Undergraduate Research Opportunities Program. Figures 1A, 1C, 2A, 2D, 2E, 6A, 6B, 6E, and S4 were created with BioRender.com. We thank Arisdelsy Cervantes, Yinze Wu, Favian Liu, and Mariya Atanasova for helpful discussions.

## References

1. DiRusso, C. J., Dashtiahangar, M., and Gilmore, T. D. (2022) Scaffold proteins as dynamic integrators of biological processes. J. Biol. Chem. 298, 102628

2. Ghosh, S. (2006) Handbook of NF-kappaB. CRC Press, Boca Rotan, FL, USA, 232 pp

3. Israël, A. (2010) The IKK complex, a central regulator of NF-κB activation. Cold Spring Harb. Perspect. Biol. 2, a000158

4. Trares, K., Ackermann, J., and Koch, I. (2022) The canonical and non-canonical NF-κB pathways and their crosstalk: A comparative study based on Petri nets. Biosystems 211, 104564

5. Maubach, G., Schmädicke, A.-C., and Naumann, M. (2017) NEMO links Nuclear Factor-κB to human diseases. Trends Mol. Med. 23, 1138–1155

6. Hanson, E. P., Monaco-Shawver, L., Solt, L. A., Madge, L. A., Banerjee, P. P., May, M. J., and Orange, J. S. (2008) Hypomorphic nuclear factor-kappaB essential modulator mutation database and reconstitution system identifies phenotypic and immunologic diversity. J. Allergy Clin. Immunol. 122, 1169–1177.e16

7. Barczewski, A. H., Ragusa, M. J., Mierke, D. F., and Pellegrini, M. (2019) The IKK-binding domain of NEMO is an irregular coiled coil with a dynamic binding interface. Sci. Rep. 9, 2950

8. Rahighi, S., Ikeda, F., Kawasaki, M., Akutsu, M., Suzuki, N., Kato, R., Kensche, T., Uejima, T., Bloor, S., Komander, D., Randow, F., Wakatsuki, S., and Dikic, I. (2009) Specific recognition of linear ubiquitin chains by NEMO is important for NF-κB activation. Cell 136, 1098–1109

9. Cordier, F., Grubisha, O., Traincard, F., Véron, M., Delepierre, M., and Agou, F. (2009) The zinc finger of NEMO is a functional ubiquitin-binding domain. J. Biol. Chem. 284, 2902–2907

10. Hinz, M., and Scheidereit, C. (2014) The IκB kinase complex in NF-κB regulation and beyond. EMBO Rep. 15, 46–61

11. Babaei, M., Liu, Y., Wuerzberger-Davis, S. M., McCaslin, E. Z., DiRusso, C. J., Yeo, A. T., Kagermazova, L., Miyamoto, S., and Gilmore, T. D. (2019) CRISPR/Cas9-based editing of a sensitive transcriptional regulatory element to achieve cell type-specific knockdown of the NEMO scaffold protein. PLoS One 14, e0222588

12. Hauenstein, A. V., Xu, G., Kabaleeswaran, V., and Wu, H. (2017) Evidence for M1-Linked polyubiquitin-mediated conformational change in NEMO. J. Mol. Biol. 429, 3793–3800

13. Shaffer, R., DeMaria, A. M., Kagermazova, L., Liu, Y., Babaei, M., Caban-Penix, S., Cervantes, A., Jehle, S., Makowski, L., Gilmore, T. D., Whitty, A., and Allen, K. N. (2019) A central region of NF-κB essential modulator is required for IKKβ-induced conformational change and for signal propagation. Biochemistry 58, 2906–2920

14. Ko, M. S., Cohen, S. N., Polley, S., Mahata, S. K., Biswas, T., Huxford, T., and Ghosh, G. (2022) Regulatory subunit NEMO promotes polyubiquitin-dependent induction of NF-κB through a targetable second interaction with upstream activator IKK2. J. Biol. Chem. 298, 101864

15. Ko, M. S., Biswas, T., Mulero, M. C., Bobkov, A. A., Ghosh, G., and Huxford, T. (2020) Structurally plastic NEMO and oligomerization prone IKK2 subunits define the behavior of human IKK2:NEMO complexes in solution. Biochim. Biophys. Acta Proteins Proteom. 1868, 140526

16. Catici, D. A. M., Amos, H. E., Yang, Y., van den Elsen, J. M. H., and Pudney, C. R. (2016) The red edge excitation shift phenomenon can be used to unmask protein structural ensembles: implications for NEMO-ubiquitin interactions. FEBS J. 283, 2272–2284

17. Catici, D. A. M., Horne, J. E., Cooper, G. E., and Pudney, C. R. (2015) Polyubiquitin drives the molecular interactions of the NF-κB essential modulator (NEMO) by allosteric regulation. J. Biol. Chem. 290, 14130–14139

18. Du, M., Ea, C.-K., Fang, Y., and Chen, Z. J. (2022) Liquid phase separation of NEMO induced by polyubiquitin chains activates NF-κB. Mol. Cell 82, 2415–2426.e5

19. Goel, S., Oliva, R., Jeganathan, S., Bader, V., Krause, L. J., Kriegler, S., Stender, I. D., Christine, C. W., Nakamura, K., Hoffmann, J.-E., Winter, R., Tatzelt, J., and Winklhofer, K. F. (2023) Linear ubiquitination induces NEMO phase separation to activate NF-κB signaling. Life Sci. Alliance. 6, e202201607

20. Sebban-Benin, H., Pescatore, A., Fusco, F., Pascuale, V., Gautheron, J., Yamaoka, S., Moncla, A., Ursini, M. V., and Courtois, G. (2007) Identification of TRAF6-dependent NEMO polyubiquitination sites through analysis of a new NEMO mutation causing incontinentia pigmenti. Hum. Mol. Genet. 16, 2805–2815

21. Gautheron, J., Pescatore, A., Fusco, F., Esposito, E., Yamaoka, S., Agou, F., Ursini, M. V., and Courtois, G. (2010) Identification of a new NEMO/TRAF6 interface affected in incontinentia pigmenti pathology. Hum. Mol. Genet. 19, 3138–3149

22. Pupko, T., Bell, R. E., Mayrose, I., Glaser, F., and Ben-Tal, N. (2002) Rate4Site: an algorithmic tool for the identification of functional regions in proteins by surface mapping of evolutionary determinants within their homologues. Bioinformatics 18 Suppl 1, S71–77

23. Golden, M. S., Cote, S. M., Sayeg, M., Zerbe, B. S., Villar, E. A., Beglov, D., Sazinsky, S. L., Georgiadis, R. M., Vajda, S., Kozakov, D., and Whitty, A. (2013) Comprehensive experimental and computational analysis of binding energy hot spots at the NF-κB essential modulator/IKKβ protein-protein interface. J. Am. Chem. Soc. 135, 6242–6256

24. Rahighi, S., Iyer, M., Oveisi, H., Nasser, S., and Duong, V. (2022) Structural basis for the simultaneous recognition of NEMO and acceptor ubiquitin by the HOIP NZF1 domain. Sci. Rep. 12, 12241

25. Dubreuil, B., and Levy, E. D. (2021) Abundance imparts evolutionary constraints of similar magnitude on the buried, surface, and disordered regions of proteins. Front. Mol. Biosci. 8, 626729

26. Schwamborn, K., Weil, R., Courtois, G., Whiteside, S. T., and Israël, A. (2000) Phorbol esters and cytokines regulate the expression of the NEMO-related protein, a molecule involved in a NF-κB-independent pathway. J. Biol. Chem. 275, 22780–22789

27. Herscovitch, M., Comb, W., Ennis, T., Coleman, K., Yong, S., Armstead, B., Kalaitzidis, D., Chandani, S., and Gilmore, T. D. (2008) Intermolecular disulfide bond formation in the NEMO dimer requires Cys54 and Cys347. Biochem. Biophys. Res. Commun. 367, 103–108

28. Cote, S. M., Gilmore, T. D., Shaffer, R., Weber, U., Bollam, R., Golden, M. S., Glover, K., Herscovitch, M., Ennis, T., Allen, K. N., and Whitty, A. (2013) Mutation of nonessential cysteines shows that the NF-κB essential modulator forms a constitutive noncovalent dimer that binds IκB kinase-β with high affinity. Biochemistry 52, 9141–9154

29. Zhou, L., Yeo, A. T., Ballarano, C., Weber, U., Allen, K. N., Gilmore, T. D., and Whitty, A. (2014) Disulfide-mediated stabilization of the IκB kinase binding domain of NF-κB essential modulator (NEMO). Biochemistry 53, 7929–7944

30. Williams, L. M., Fuess, L. E., Brennan, J. J., Mansfield, K. M., Salas-Rodriguez, E., Welsh, J., Awtry, J., Banic, S., Chacko, C., Chezian, A., Dowers, D., Estrada, F., Hsieh, Y.-H., Kang, J., Li, W., Malchiodi, Z., Malinowski, J., Matuszak, S., McTigue, T., Mueller, D., Nguyen, B., Nguyen, M., Nguyen, P., Nguyen, S., Njoku, N., Patel, K., Pellegrini, W., Pliakas, T., Qadir, D., Ryan, E., Schiffer, A., Thiel, A., Yunes, S. A., Spilios, K. E., Pinzón C, J. H., Mydlarz, L. D., and Gilmore, T. D. (2018) A conserved Toll-like receptor-to-NF-κB signaling pathway in the endangered coral *Orbicella faveolata*. Dev. Comp. Immunol. 79, 128–136

31. Scholefield, J., Henriques, R., Savulescu, A. F., Fontan, E., Boucharlat, A., Laplantine, E., Smahi, A., Israël, A., Agou, F., and Mhlanga, M. M. (2016) Super-resolution microscopy reveals a preformed NEMO lattice structure that is collapsed in incontinentia pigmenti. Nat. Commun. 7, 12629

32. Tarantino, N., Tinevez, J.-Y., Crowell, E. F., Boisson, B., Henriques, R., Mhlanga, M., Agou, F., Israël, A., and Laplantine, E. (2014) TNF and IL-1 exhibit distinct ubiquitin requirements for inducing NEMO-IKK supramolecular structures. J. Cell Biol. 204, 231–245

33. Tian, W., Zhao, C., Hu, Q., Sun, J., and Peng, X. (2016) Roles of Toll-like receptors 2 and 6 in the inflammatory response to *Mycoplasma gallisepticum* infection in DF-1 cells and in chicken embryos. Dev. Comp. Immunol. 59, 39–47

34. Parra-Izquierdo, I., Lakshmanan, H. H. S., Melrose, A. R., Pang, J., Zheng, T. J., Jordan, K. R., Reitsma, S. E., McCarty, O. J. T., and Aslan, J. E. (2021) The Toll-Like Receptor 2 ligand Pam2CSK4 activates platelet nuclear factor-κB and Bruton’s Tyrosine Kinase signaling to promote platelet-endothelial cell interactions. Front. Immunol. 12, 729951

35. Rushe, M., Silvian, L., Bixler, S., Chen, L. L., Cheung, A., Bowes, S., Cuervo, H., Berkowitz, S., Zheng, T., Guckian, K., Pellegrini, M., and Lugovskoy, A. (2008) Structure of a NEMO/IKK-associating domain reveals architecture of the interaction site. Structure 16, 798–808

36. Lo, Y.-C., Lin, S.-C., Rospigliosi, C. C., Conze, D. B., Wu, C.-J., Ashwell, J. D., Eliezer, D., and Wu, H. (2009) Structural basis for recognition of diubiquitins by NEMO. Mol. Cell 33, 602–615

37. Bagnéris, C., Ageichik, A. V., Cronin, N., Wallace, B., Collins, M., Boshoff, C., Waksman, G., and Barrett, T. (2008) Crystal structure of a vFlip-IKKγ complex: insights into viral activation of the IKK signalosome. Mol. Cell 30, 620–631

38. Pace, C. N. (1990) Measuring and increasing protein stability. Trends Biotechnol. 8, 93–98

39. Fujino, Y., Miyagawa, T., Torii, M., Inoue, M., Fujii, Y., Okanishi, H., Kanai, Y., and Masui, R. (2021) Structural changes induced by ligand binding drastically increase the thermostability of the Ser/Thr protein kinase TpkD from *Thermus thermophilus* HB8. FEBS Lett. 595, 264–274

40. Guo, B., Audu, C. O., Cochran, J. C., Mierke, D. F., and Pellegrini, M. (2014) Protein engineering of the N-terminus of NEMO: structure stabilization and rescue of IKKβ binding. Biochemistry 53, 6776–6785

41. Mueterthies, J., and Potoyan, D. A. (2021) Solvent exposure and ionic condensation drive fuzzy dimerization of disordered heterochromatin protein sequence. Biomolecules. 11, 915

42. Jencks, W. P. (1975) Binding energy, specificity, and enzymic catalysis: the circe effect. Adv. Enzymol. Relat. Areas Mol. Biol. 43, 219–410

43. Whitty, A., Fierke, C. A., and Jencks, W. P. (1995) Role of binding energy with coenzyme A in catalysis by 3-oxoacid coenzyme A transferase. Biochemistry 34, 11678–11689

44. Zhu, G., Wu, C.-J., Zhao, Y., and Ashwell, J. D. (2007) Optineurin negatively regulates TNFα-induced NF-κB activation by competing with NEMO for ubiquitinated RIP. Curr. Biol. 17, 1438–1443

45. Gough, N. R. (2007) Optineurin competes with NEMO. Science’s STKE 2007, tw310–tw310

46. Maruyama, H., Morino, H., Ito, H., Izumi, Y., Kato, H., Watanabe, Y., Kinoshita, Y., Kamada, M., Nodera, H., Suzuki, H., Komure, O., Matsuura, S., Kobatake, K., Morimoto, N., Abe, K., Suzuki, N., Aoki, M., Kawata, A., Hirai, T., Kato, T., Ogasawara, K., Hirano, A., Takumi, T., Kusaka, H., Hagiwara, K., Kaji, R., and Kawakami, H. (2010) Mutations of optineurin in amyotrophic lateral sclerosis. Nature 465, 223–226

47. Pakravan, D., Orlando, G., Bercier, V., and Van Den Bosch, L. (2021) Role and therapeutic potential of liquid-liquid phase separation in amyotrophic lateral sclerosis. J. Mol. Cell. Biol. 13, 15–28

48. Rothwarf, D. M., Zandi, E., Natoli, G., and Karin, M. (1998) IKK-γ is an essential regulatory subunit of the IκB kinase complex. Nature 395, 297–300

49. Clark, K., Nanda, S., and Cohen, P. (2013) Molecular control of the NEMO family of ubiquitin-binding proteins. Nat. Rev. Mol. Cell. Biol. 14, 673–685

50. Schröfelbauer, B., Polley, S., Behar, M., Ghosh, G., and Hoffmann, A. (2012) NEMO ensures signaling specificity of the pleiotropic IKKβ by directing its kinase activity toward IκBα. Mol. Cell 47, 111–121

51. Lee, S. H., Toth, Z., Wong, L.-Y., Brulois, K., Nguyen, J., Lee, J.-Y., Zandi, E., and Jung, J. U. (2012) Novel phosphorylations of IKKγ/NEMO. mBio 3, e00411–12

52. Jackson, S. S., Coughlin, E. E., Coon, J. J., and Miyamoto, S. (2013) Identifying post-translational modifications of NEMO by tandem mass spectrometry after high affinity purification. Protein Expr. Purif. 92, 48–53

53. Yoshizawa, T., Nozawa, R.-S., Jia, T. Z., Saio, T., and Mori, E. (2020) Biological phase separation: cell biology meets biophysics. Biophys. Rev. 12, 519–539

54. Ensinck, M. M., and Carlon, M. S. (2022) One size does not fit all: the past, present and future of cystic fibrosis causal therapies. Cells 11, 1868

55. Sievers, F., Wilm, A., Dineen, D., Gibson, T. J., Karplus, K., Li, W., Lopez, R., McWilliam, H., Remmert, M., Söding, J., Thompson, J. D., and Higgins, D. G. (2011) Fast, scalable generation of high-quality protein multiple sequence alignments using Clustal Omega. Mol. Syst. Biol. 7, 539

56. Brown, N. P., Leroy, C., and Sander, C. (1998) MView: a web-compatible database search or multiple alignment viewer. Bioinformatics 14, 380–381

57. Shannon, P., Markiel, A., Ozier, O., Baliga, N. S., Wang, J. T., Ramage, D., Amin, N., Schwikowski, B., and Ideker, T. (2003) Cytoscape: a software environment for integrated models of biomolecular interaction networks. Genome Res. 13, 2498–2504

58. Wolenski, F. S., Garbati, M. R., Lubinski, T. J., Traylor-Knowles, N., Dresselhaus, E., Stefanik, D. J., Goucher, H., Finnerty, J. R., and Gilmore, T. D. (2011) Characterization of the core elements of the NF-κB signaling pathway of the sea anemone *Nematostella vectensis*. Mol. Cell. Biol. 31, 1076– 1087

59. Gasteiger, E., Hoogland, C., Gattiker, A., Duvaud, S.. Wilkins, M. R., Appel, R. D., Bairoch, A. (2005) Protein identification and analysis tools on the ExPASy server. In The Proteomics Protocols Handbook; Walker, J. M., Ed.; Humana Press: Totowa, NJ, pp 571–607

60. Simon, M. D., Heider, P. L., Adamo, A., Vinogradov, A. A., Mong, S. K., Li, X., Berger, T., Policarpo, R. L., Zhang, C., Zou, Y., Liao, X., Spokoyny, A. M., Jensen, K. F., and Pentelute, B. L. (2014) Rapid flow-based peptide synthesis. ChemBioChem 15, 713–720

61. Garbati, M. R., Alço, G., and Gilmore, T. D. (2010) Histone acetyltransferase p300 is a coactivator for transcription factor REL and is C-terminally truncated in the human diffuse large B-cell lymphoma cell line RC-K8. Cancer Lett. 291, 237–245

62. Greenfield, N. J. (2006) Using circular dichroism spectra to estimate protein secondary structure. Nat. Protoc. 1, 2876–2890

63. Williams, L. M., Inge, M. M., Mansfield, K. M., Rasmussen, A., Afghani, J., Agrba, M., Albert, C., Andersson, C., Babaei, M., Babaei, M., Bagdasaryants, A., Bonilla, A., Browne, A., Carpenter, S., Chen, T., Christie, B., Cyr, A., Dam, K., Dulock, N., Erdene, G., Esau, L., Esonwune, S., Hanchate, A., Huang, X., Jennings, T., Kasabwala, A., Kehoe, L., Kobayashi, R., Lee, M., LeVan, A., Liu, Y., Murphy, E., Nambiar, A., Olive, M., Patel, D., Pavesi, F., Petty, C. A., Samofalova, Y., Sanchez, S., Stejskal, C., Tang, Y., Yapo, A., Cleary, J. P., Yunes, S. A., Siggers, T., and Gilmore, T. D. (2020) Transcription factor NF-κB in a basal metazoan, the sponge, has conserved and unique sequences, activities, and regulation. Dev. Comp. Immunol. 104, 103559

