## Supporting Information for "A Conserved Core Region of the Scaffold NEMO is Essential for Signal-induced Conformational Change and Liquid-liquid Phase Separation"

***Corresponding authors**

**This file includes:**

**Figures 1 to 8**

**Tables 1 and 2**

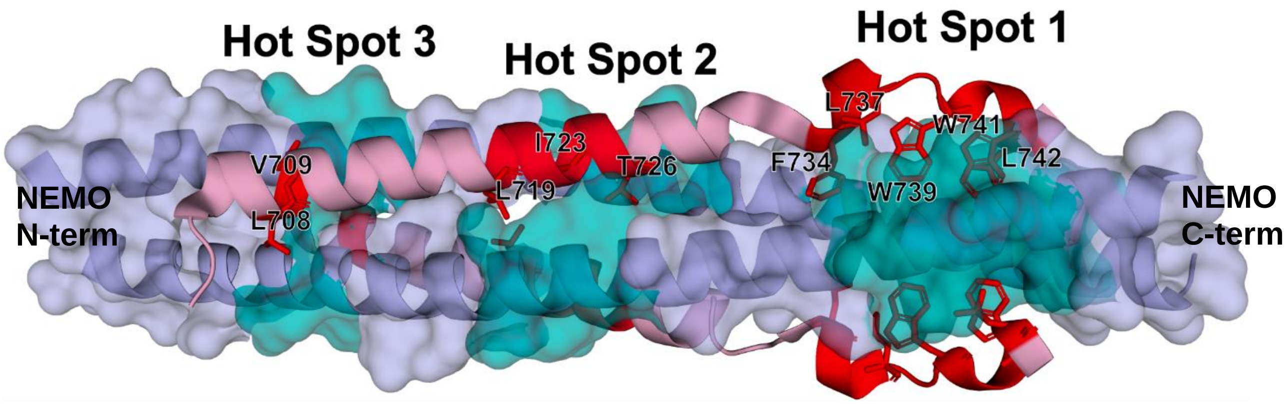


**Figure S1. NEMO/IKKβ interaction hot spots.** The crystal structure of the NEMO/IKK$\beta$ interface (PDB ID 3brv). IKK$\beta$ is in red, with the residues found to be involved in hot spot interactions highlighted in bright red and labelled. NEMO is in blue, with regions involved in the hot spot interactions highlighted in teal.

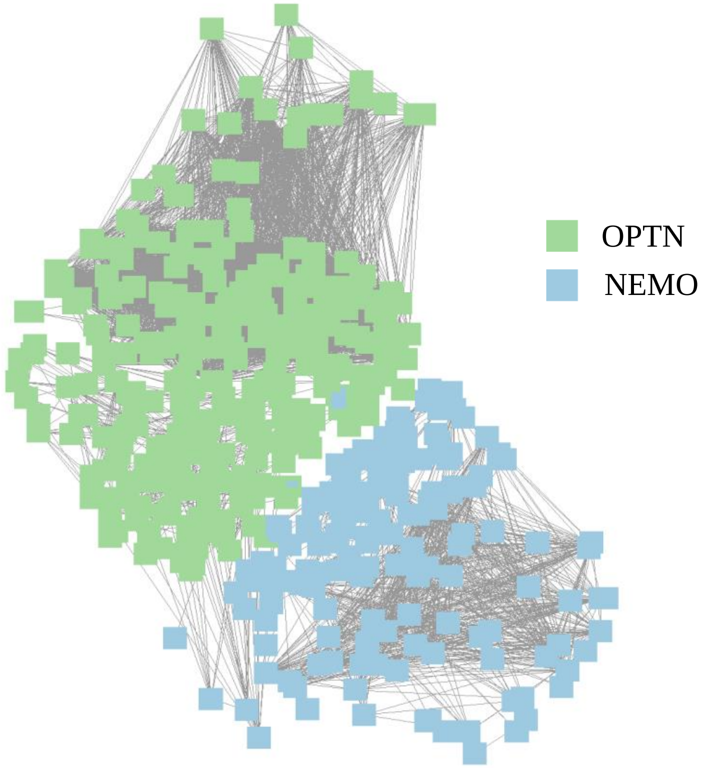


**Figure S2. Sequence Similarity Network (SSN) of NEMO and optineurin (OPTN).** SSN representation of a BLAST sequence search conducted using human NEMO with a UniProt query e-value of 1 x $10^{-5}$, with an alignment score cut-off of 61%, with 674 retrieved sequences and 2708 nodes. Network was visualized using Cytoscape.

**Figure S3. NEMO and OPTN multiple sequence alignment. (Page 1)**

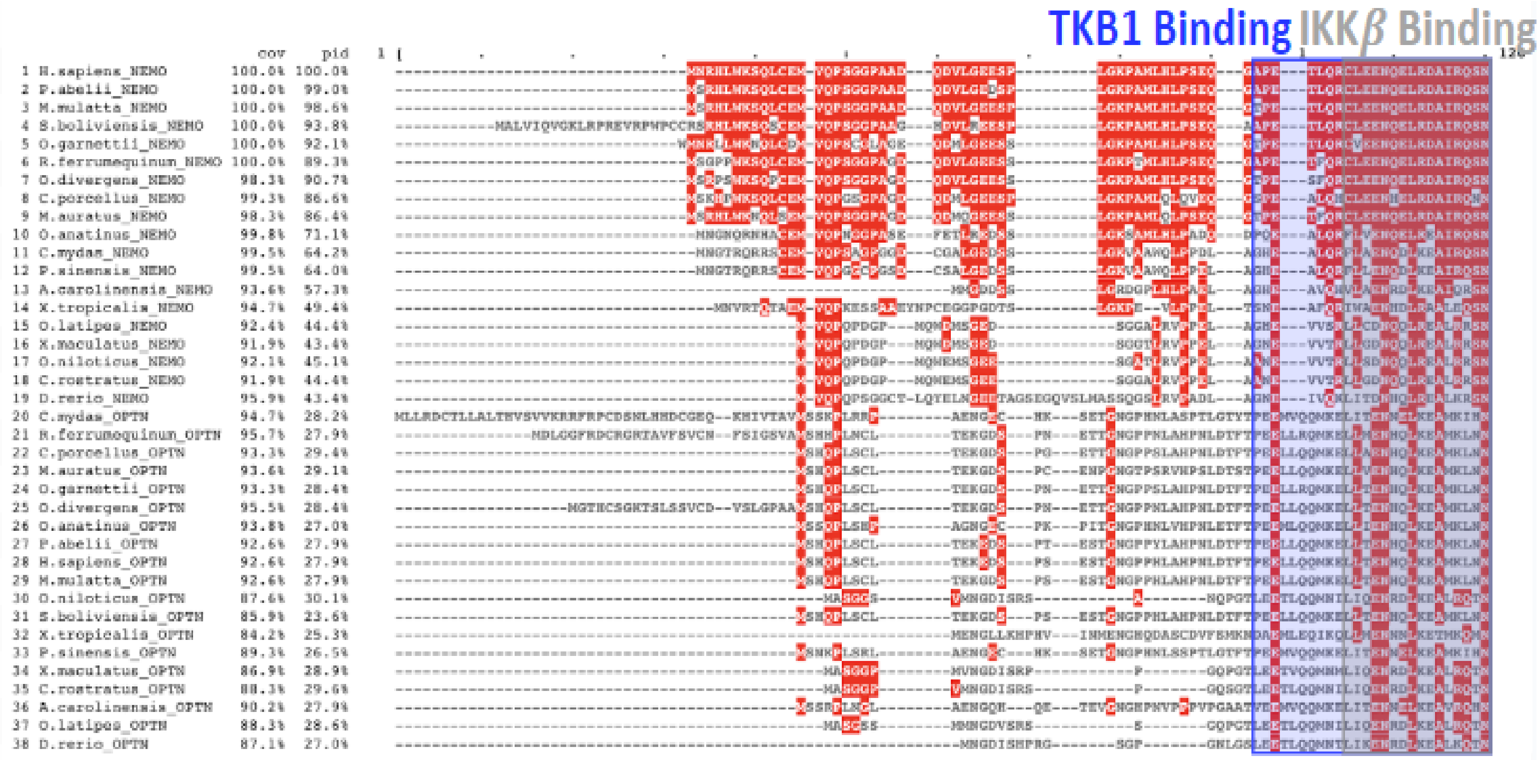


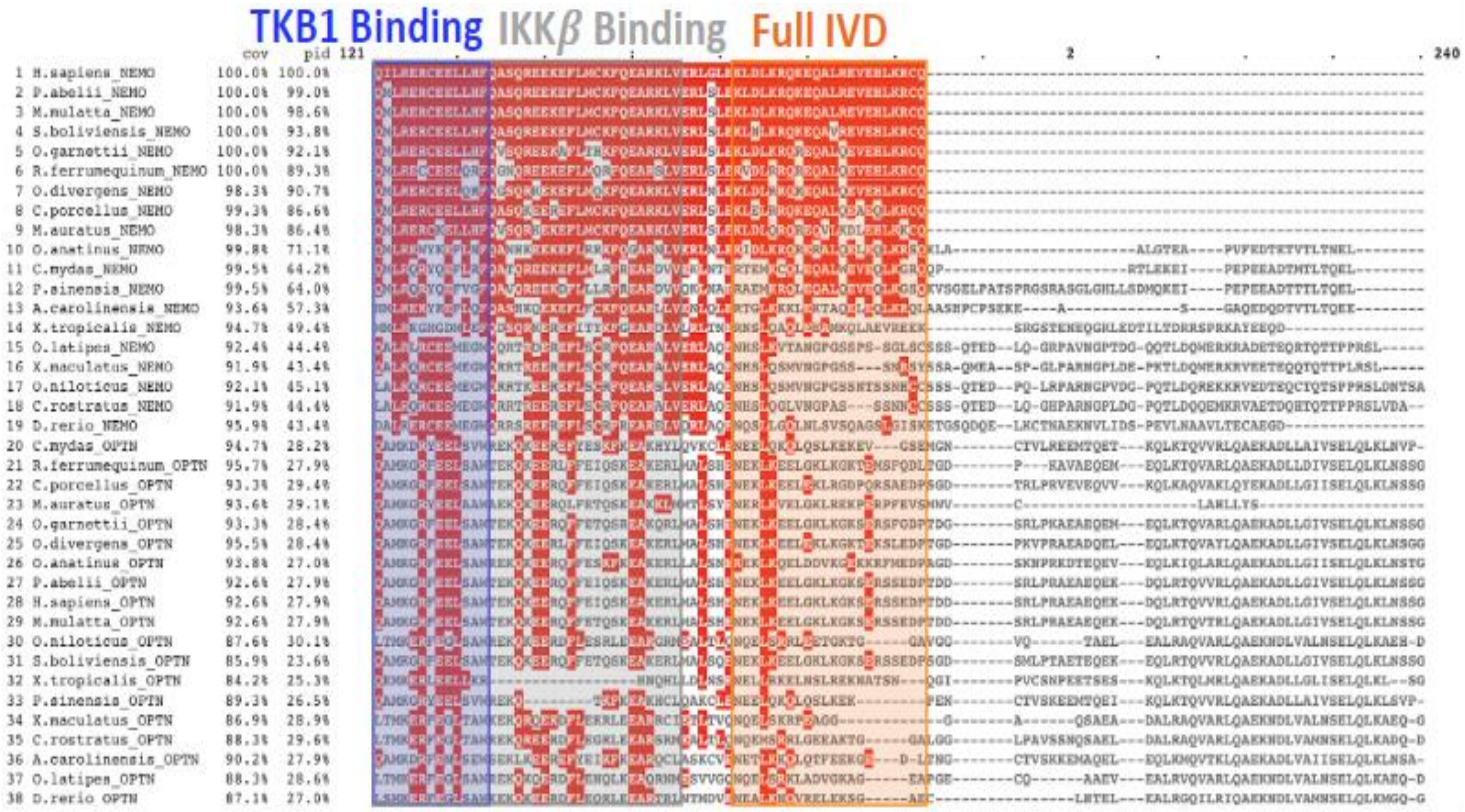

**Figure S3. NEMO and OPTN multiple sequence alignment. (Page 2)**

```
                            cov    pid 241          :         .         .         .         .         3         .         .         .         .         :         . 360
 1 H.sapiens_NEMO         100.0% 100.0%    ------------------------------------------------------------------------------------------------------------------------
 2 P.abelii_NEMO          100.0%  99.0%    ------------------------------------------------------------------------------------------------------------------------
 3 M.mulatta_NEMO         100.0%  98.6%    ------------------------------------------------------------------------------------------------------------------------
 4 S.boliviensis_NEMO     100.0%  93.8%    ------------------------------------------------------------------------------------------------------------------------
 5 O.garnettii_NEMO       100.0%  92.1%    ------------------------------------------------------------------------------------------------------------------------
 6 R.ferrumequinum_NEMO   100.0%  89.3%    ------------------------------------------------------------------------------------------------------------------------
 7 O.divergens_NEMO        98.3%  90.7%    ------------------------------------------------------------------------------------------------------------------------
 8 C.porcellus_NEMO        99.3%  86.6%    ------------------------------------------------------------------------------------------------------------------------
 9 M.auratus_NEMO          98.3%  86.4%    ------------------------------------------------------------------------------------------------------------------------
10 O.anatinus_NEMO         99.8%  71.1%    --------VDLA----------------MVPGEKTDYLQALKEQKEQLENHLQQLQKTNHSLLMEKLALLTEN-----QELKLKVPPRQPEEPRLKVKENGILPERLLETESLQGASGS-
11 C.mydas_NEMO            99.5%  64.2%    --------AEPP---------------MMAPGSETLQLL--KEQ-------LEQLQEANQTL-----------------QAQIQQQVQPPGEAGVRVKENGVQLERPVELEALQGSGDSG
12 P.sinensis_NEMO         99.5%  64.0%    --------AEPT----------------MAPGQETQQLE--QH------------QEA----------------------SQVQQQVQPS-NVGL-KQENGVQLERPVELETLQGSGDSR
13 A.carolinensis_NEMO     93.6%  57.3%    --------NIFA----------------VVPES-------TQQNRGQLEEELKVLQDANQALQKEKLALQAEI-----CA------LQQATKTTLPVEENGLQAERPLELESLQGSGDRK
14 X.tropicalis_NEMO       94.7%  49.4%    ---------------------------YATPGSSLSSCE----------------------------TLTHSI-----CETIRQTVPDEEGPSRNKQKENGN-KEVCMENEIMQGTVDRN
15 O.latipes_NEMO          92.4%  44.4%    -----------------------------PLEGTNDFLKLLKSHKEQLEETMRDLMKKNENLEREKEEGEKERDRMRRCIDQLRSKLLH--TQATS-----------GTEEVVQLRSEAQ
16 X.maculatus_NEMO        91.9%  43.4%    -----------------------------PAEGANEFLQLLKSHKEKTEEEMREMRRKNEELKRERDEGKEVLERQHRCIEHLRAKLFH--TQMSS------------NEEAVQLRSEAQ
17 O.niloticus_NEMO        92.1%  45.1%    HSSLSYAQSSPPISFSDELGDILGNDNSQPVEGANEFLQLLKSHKEKLEEGMRELRRKNEELEKQRDEGEKEKEQMLHCIDQLRGKLAQ--AQTNN-----------VTEEVVQQRSEAQ
18 C.rostratus_NEMO        91.9%  44.4%    NSSLCSALPSPPTSLSEVGGEIMSRENGQPVEGANEFLQLLKSHKEKLEEGMRELRRKNDELERERDEGEKERERMRRCIDQLRAKLAQ--AQANS-----------VTEEVVQHRSEAQ
19 D.rerio_NEMO            95.9%  43.4%    -------------------TDRHTMPQSLPAVGSNDFLKLLKSHKEKLEEGMRALRRKNEELEKEKAENEKERENLLAALEQLRSKLSQ--NVG-------------VTEESVQP-----
20 C.mydas_OPTN            94.7%  28.2%    S-------------------------------TEDSFVEIRMSEGE-PNAALKENQDVVN--------------------DQRTNNAAA-YIRSRSVDEPK-NLESE-ELTVSQLLC--C
21 R.ferrumequinum_OPTN    95.7%  27.9%    H-------------------------------SEDSFVEIRMAEGE-AHVAVKEMKTSPG----------------PTRTDFIDT--------SRSAESARDYLESE-ELTVSQLLL--C
22 C.porcellus_OPTN        93.3%  29.4%    S-------------------------------CEDSFVEIRMAEGE-ADGAVKEIKSSCP---------------RLPRTDSTDM--------SKSVEGTKKYLEFELELTVSQLLL--C
23 M.auratus_OPTN          93.6%  29.1%    -----------------------------------VAFVSALQEGD-AEGAMKEMKSSSI----------------PVRTDAINM--------TKSTEDTRTCVESE-ELAVSQLLP--C
24 O.garnettii_OPTN        93.3%  28.4%    P-------------------------------TEDSFVEIRMAEGE-ADGSVKEIKNSSG----------------PIRTDSTVTFLSS-YIRSRSEEGAKNYLEFE-ELTVSQLLL--C
25 O.divergens_OPTN        95.5%  28.4%    P-------------------------------SEDSFVEIRIAEGE-ADVAVREIRTSPG----------------TARIDSTDT--------NKSTERSRNYLEFE-KLTVSQLLL--C
26 O.anatinus_OPTN         93.8%  27.0%    S-------------------------------SEDSFVEIRMAEGD-ETRDLKETMNGSQ----------------PRPADN--------FDTSKIVNAAKSYSERE-ELTVSQLLL--C
27 P.abelii_OPTN           92.6%  27.9%    S-------------------------------SEDSFVEIRMAEGE-AEGSVKEIKHSPG----------------PTRTVST-------SR-SRSAEGAKNYLEHE-ELTVSQLLL--C
28 H.sapiens_OPTN          92.6%  27.9%    S-------------------------------SEDSFVEIRMAEGE-AEGSVKEIKHSPG----------------PTRTVSTGTALSK-YR-SRSADGAKNYFEHE-ELTVSQLLL--C
29 M.mulatta_OPTN          92.6%  27.9%    S-------------------------------SEDSFVEIRMAEGE-AEGSVKEIKHSPG----------------PTRTVSIGT--------SRSAEGAKNYLEHE-ELTVSQLLL--C
30 O.niloticus_OPTN        87.6%  30.1%    S-------------------------------QDDSFIEIIKVSDG-GVDGMNEALGAER----------------SR---------------RQDACMTASRMDNE-EVTVSQLLQ--S
31 S.boliviensis_OPTN      85.9%  23.6%    S-------------------------------SEDSFVEIRMAEGE-AEGSVKEIKHSPG----------------PTRKVPIGT-----YV-KEDAEGAKNYLEHE-ELTVSQLLL--C
32 X.tropicalis_OPTN       84.2%  25.3%    F-------------------------------SEDSFVEIRISEKE-SGEIIHEEKANKT---------------LPDY--------DV-SYRTNSVKEEGSGTEPE-EVAISRLLR--S
33 P.sinensis_OPTN         89.3%  26.5%    S-------------------------------TEDSFVEIRMSEGV-PNAALKENQDIAN--------------------DQRISNPAA-YIRSGSVDEPK-NLESE-EFTVRQLLC--C
34 X.maculatus_OPTN        86.9%  28.9%    A-------------------------------EEDSFIEIIKVSDA-GVEDMNQTCGSER----------------IR---------------RPDLSMTTSRQENE-EATVSRLLQ--S
35 C.rostratus_OPTN        88.3%  29.6%    S-------------------------------HSDSFIEIIRVTDG-GVDGVNDVCGLER----------------SR---------------RPDLSMTASRLDSE-EVTVSQLLQ--S
36 A.carolinensis_OPTN     90.2%  27.9%    T-------------------------------AEDSFVEIRMHEEE-VNGALKENQDLS---------------------DKRMGDTPT-YVRSRSVDEAKKYLQTE-ELTVSQLLH--S
37 O.latipes_OPTN          88.3%  28.6%    S-------------------------------RDDSFIEIIRVTDG-GVDAVNEARGAER----------------GR---------------CPELGMTASRLDNE-EATVSQLLQ--S
38 D.rerio_OPTN            87.1%  27.0%    S-------------------------------PSNSFIEIRIAE----------------------------------------------------------------------------
```

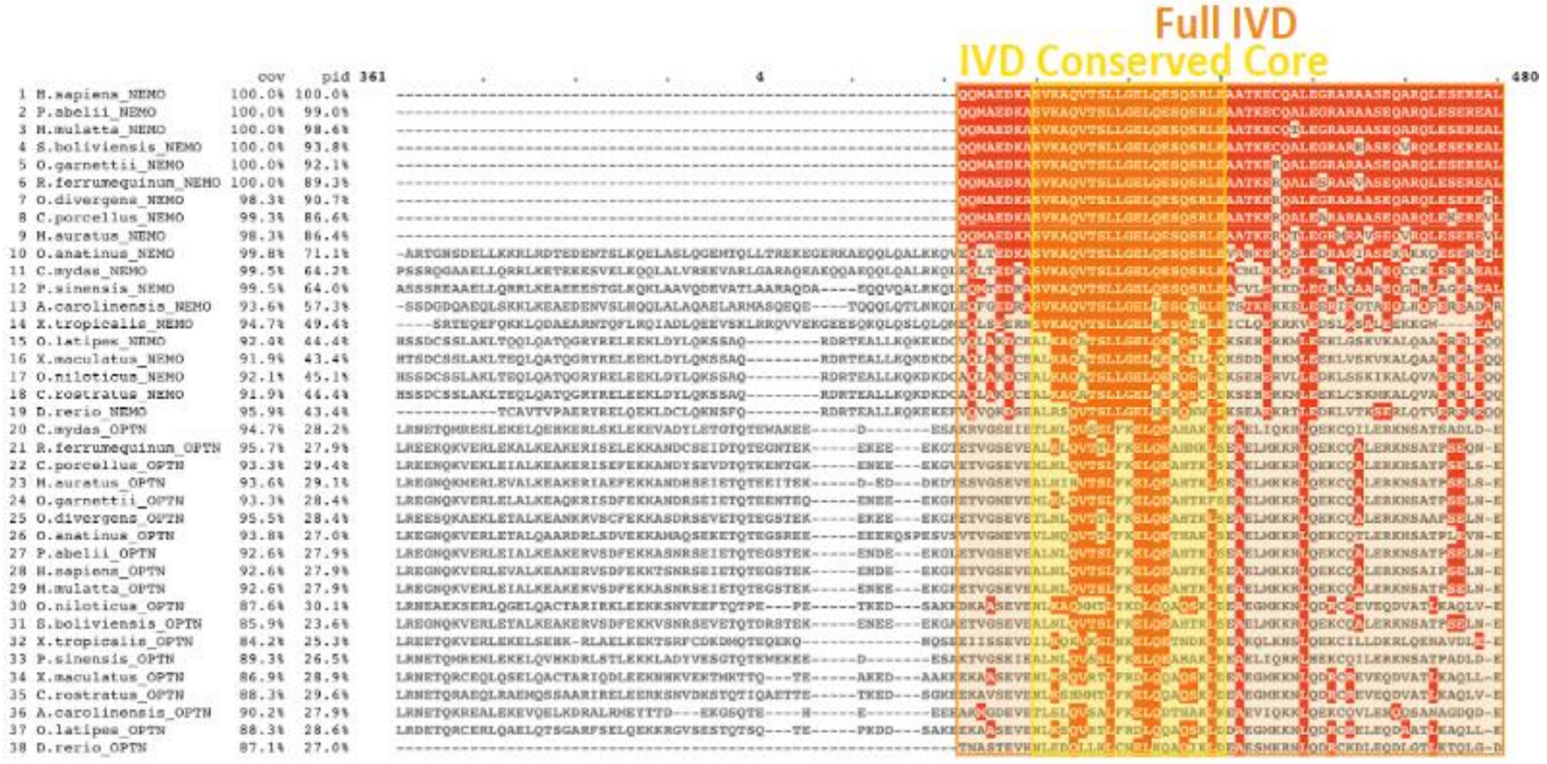

**Figure S3. NEMO and OPTN multiple sequence alignment. (Page 3)**

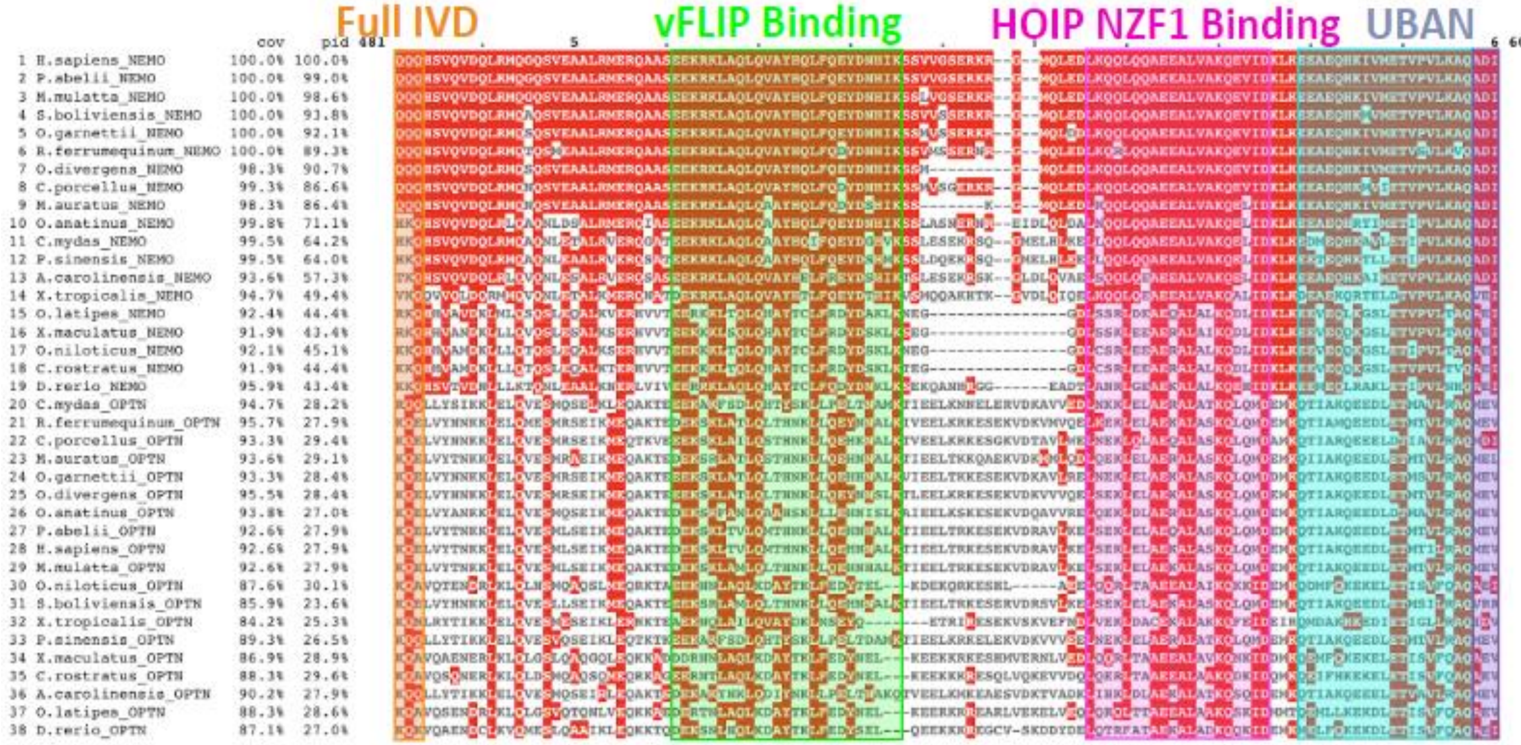


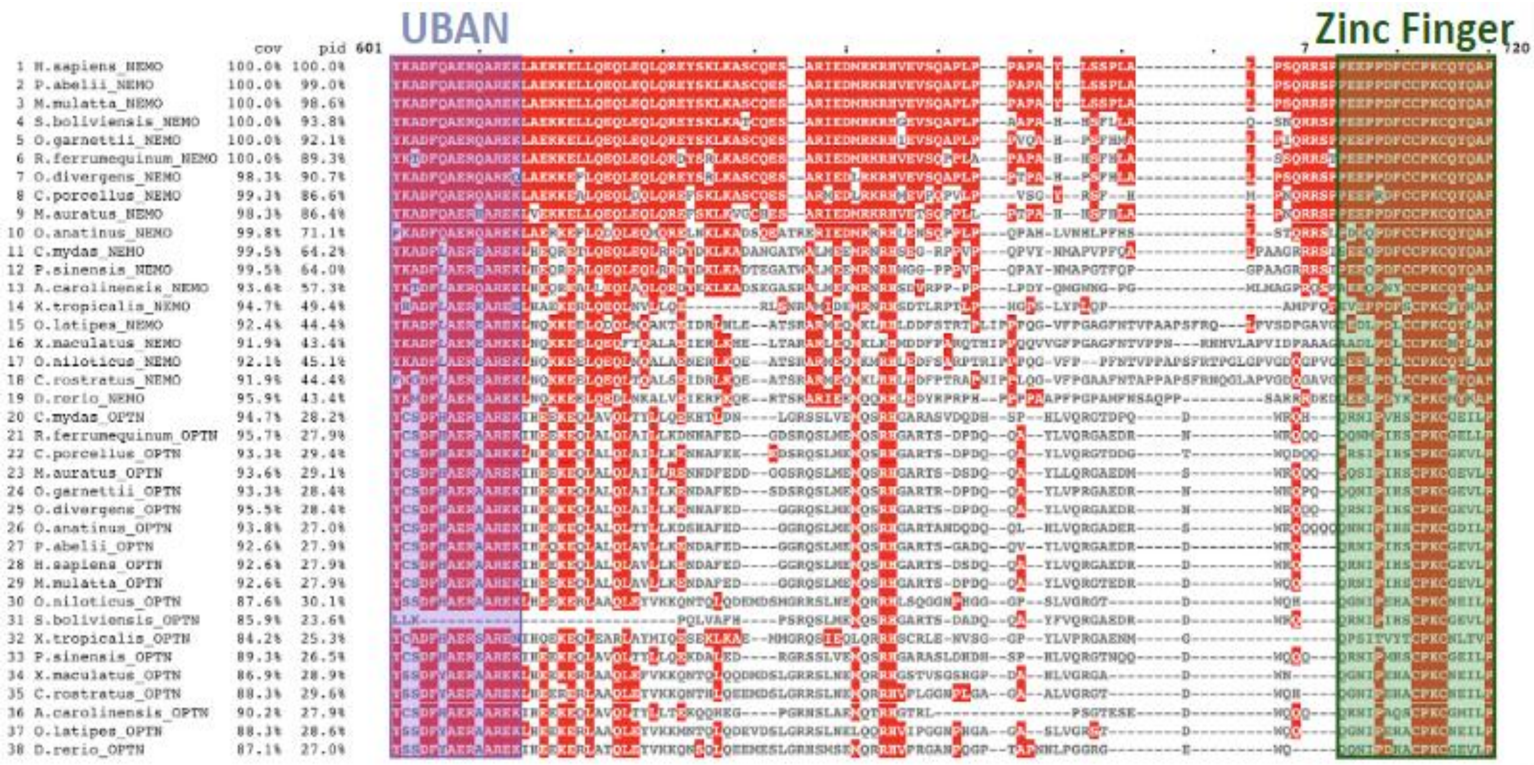

**Figure S3. NEMO and OPTN multiple sequence alignment. (Page 4)**

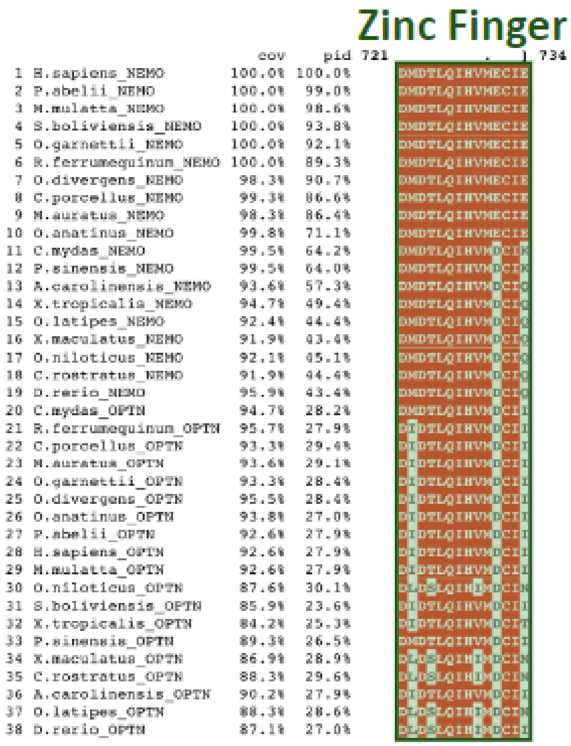


**Figure S3. NEMO and OPTN multiple sequence alignment.** Alignment of NEMO and OPTN from select species using Clustal-Omega and visualized using MView. Residues highlighted in red are identical to the corresponding residue in human NEMO.

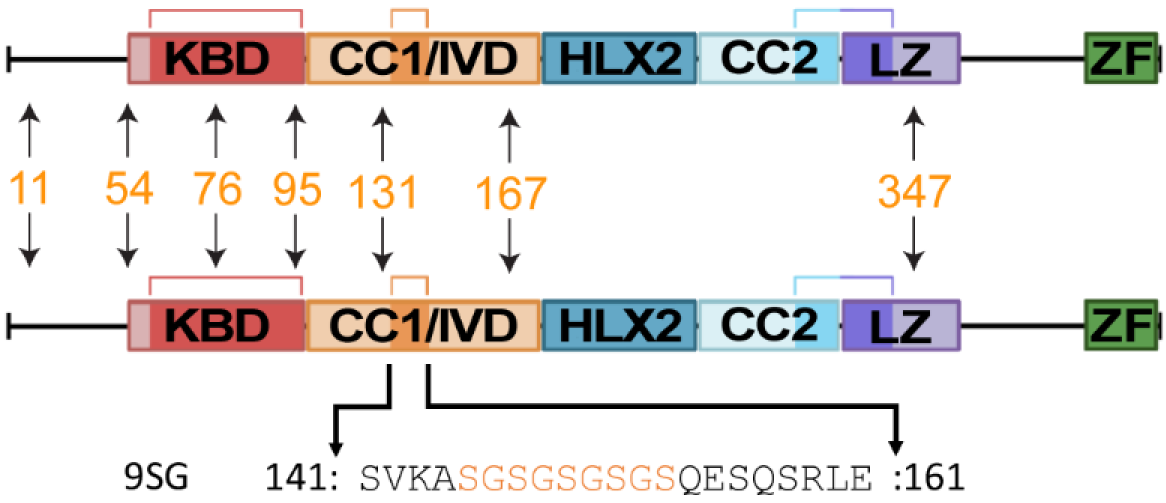


**Figure S4. Comparison of the 7XAla and 9SG-7XAla NEMO proteins.** Domain maps of 7XAla-NEMO and 9SG (7XAla background) constructs used in this study. Orange numbers represent the seven native cysteines that were changed to alanine (11, 54, 76, 95, 131, 167, 347) in both constructs. Orange letters indicate the mutation of residues 145-153 to the Ser-Gly repeat SGSGSGSGS.

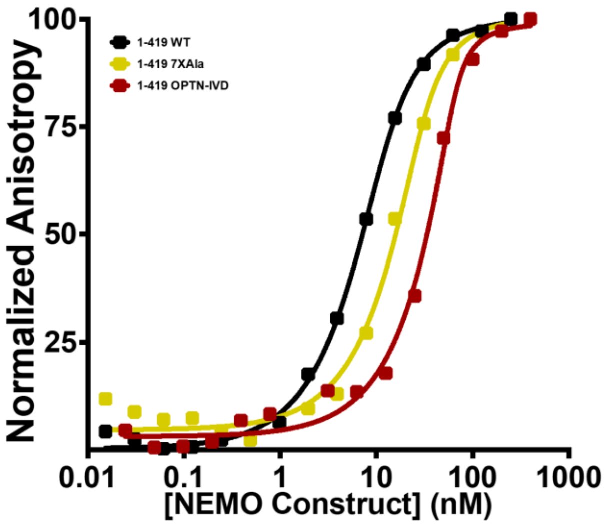


**Figure S5. IKKβ 45-mer peptide binding to full-length NEMO variants.** Fluorescence anisotropy binding assays of full-length NEMO proteins (aa 1-419). The concentrations of the NEMO proteins were varied, while the concentration of FITC-labelled IKKβ (701−745) was kept constant at 15 nM. Results represent averages from at least two independent experiments.

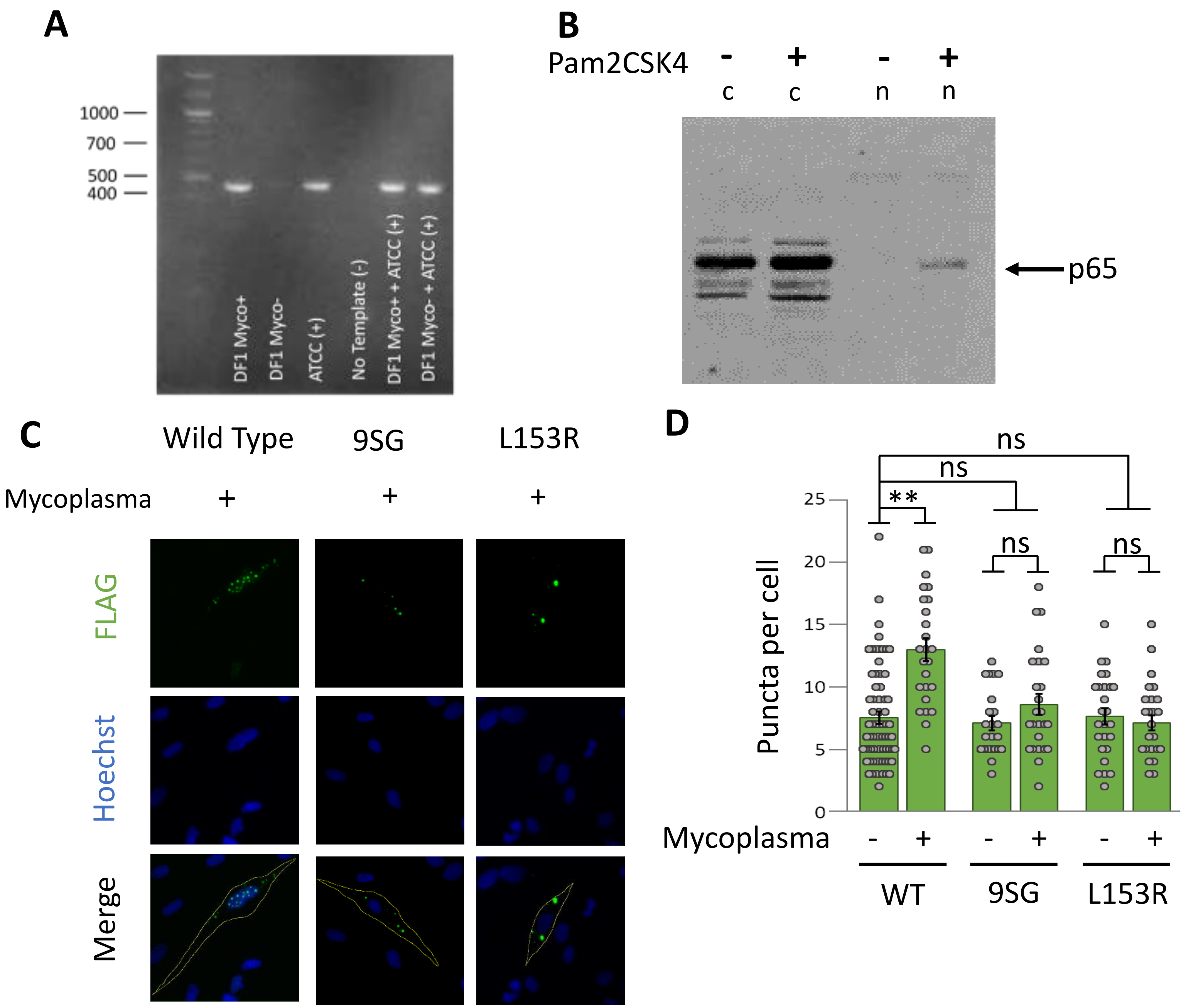


**Figure S6. Mycoplasma infection chronically activates the NF-κB pathway in DF-1 chicken fibroblasts.** (A) Agarose gel confirming the infection of DF-1 chicken fibroblasts with mycoplasma using the ATTC PCR mycoplasma detection kit. Molecular weight markers are indicated (in base pairs) on the left. (B) Anti-p65 Western blot of cytosolic (c) and nuclear (n) extracts from DF-1 chicken fibroblasts. After confirming mycoplasma infection, cells were scraped from 100-mm cell culture dishes and incubated in hypotonic buffer (10 mM HEPES pH 7.9, 1.5 mM $MgCl_2$, 10 mM KCl) for 10 min on ice, which was then supplemented with NP-40 to a concentration of 0.5%. Samples were then vortexed, and pelleted at 800 x g for 5 min at 4°C. The cytosolic fraction was collected from the supernatant, while the nuclear pellet was washed with hypotonic buffer and re-pelleted. The nuclear pellet was re-suspended in hypertonic buffer (20 mM HEPES pH 7.9, 1.5 mM $MgCl_2$, 0.2 mM EDTA, 420 mM NaCl, 25% v/v glycerol) and rocked for 1 h. The nuclear extract was clarified by centrifugation at 13,000 rpm for 30 min. (C) Representative immunofluorescence images of puncta formation in DF-1 cells transfected with the expression vectors for the indicated FLAG-tagged NEMO proteins. Cells were confirmed for mycoplasma infection as in (A) at 24 h prior to transfection using Effectene. Forty-eight h after transfection, cells were subjected to extraction with saponin buffer and then fixed with 4% paraformaldehyde and permeabilized with 0.2% Triton-X100. (D) Quantitation of puncta per cell of transfected cells in (C). The control uninfected cells were from Fig. 5C in main text. Shown are the means ± SEM; n = $\geq$ 20 cells per condition in at least two experiments. ns, not significant; **, $p<0.001$ by t-test assuming equal or unequal variance as appropriate. Dots are the number of puncta in individual cells.

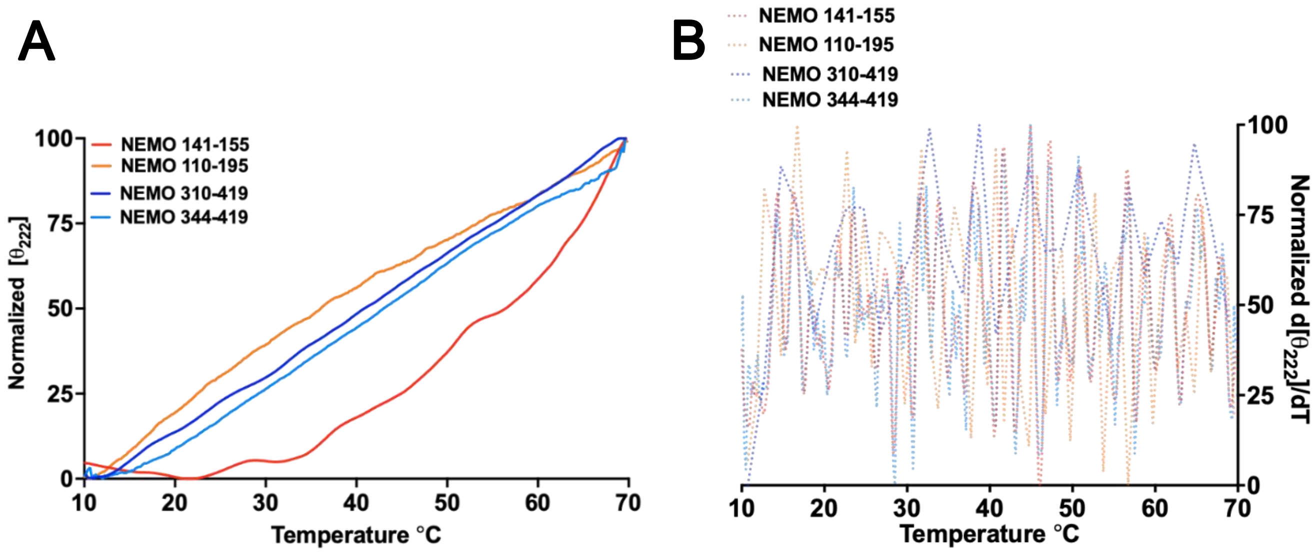


**Figure S7. Constructs displaying no visible cooperative thermal denaturation event via CD.** (A) The CD-monitored thermal denaturation measuring the loss in secondary structure of the indicated NEMO constructs as an increase in the 222 nm signal. (B) Plot showing the first derivative of the denaturation plot in (A).

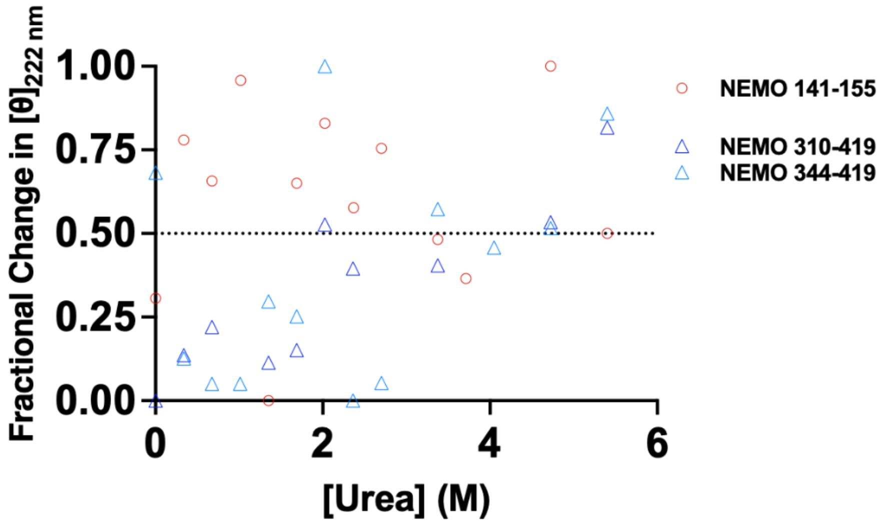


**Figure S8. Constructs displaying no visible cooperative chemical denaturation event, as monitored via CD**. Shown is the CD-monitored chemical denaturation using urea as the denaturant, measuring the loss in secondary structure of the indicated NEMO constructs as an increase in the 222 nm signal.

**Supplemental Table 1. Plasmids used in this study.**

| Plasmid Name | Description/Source |
|---|---|
| **pcDNA-FLAG** | Ref. (27) |
| **pcDNA-FLAG NEMO** | Ref. (27) |
| **pcDNA-FLAG-NEMO-9SG** | Ref. (13) |
| **pcDNA-FLAG-NEMO-L153R** | Ref. (11) |
| **pUC57-NEMO-Δ145-153** | pUC57-Simple with human NEMO cDNA containing amino acids 2-110 followed by 196-419 of human NEMO. Has a 5′ EcoRI and 3′ BamHI sites for excision. Synthesized by GenScript. |
| **pcDNA-FLAG-NEMO-Δ145-153** | EcoRI-BamHI fragment containing amino acids 2-110 followed by 196-419 of human NEMO was excised from pUC57-NEMO-Δ145-153 (GenScript) and subcloned into EcoRI-BamHI-digested pcDNA-FLAG. |
| **pUC57-OPTN-IVD-NEMO** | pUC57-Simple with human NEMO cDNA containing amino acids 2-419 of human NEMO containing the following variation at nucleotides 420-474: TGAATCTCCAGGTGACGTCCTTGTTCAAGGAGCTGCAGGAGGCCCATACTAAA. Has a 5′ EcoRI and 3′ BamHI sites for excision. Synthesized by GenScript. |
| **FLAG OPTN-IVD-NEMO** | EcoRI-BamHI fragment containing amino acids 2-419 of human NEMO was excised from pUC57-OPTN-IVD-NEMO (GenScript) and subcloned into EcoRI-BamHI-digested pcDNA-FLAG. |

| | |
|---|---|
| **pUC57-9SG-L153** | pUC57-Simple with human NEMO cDNA containing amino acids 2-419 of human NEMO containing the following variation at nucleotides 429-461: TCTGGGTCTGGGTCTGGGTCTGGG. Has a 5’ EcoRI and 3’ BamHI sites for excision. Synthesized by GenScript. |
| **FLAG 9SG-L153** | EcoRI-BamHI fragment containing amino acids 2-419 of human NEMO was excised from pUC57-9SG-L153 (GenScript) and subcloned into EcoRI-BamHI-digested pcDNA-FLAG. |
| **pE-SUMOstar Amp** | LifeSensors (PE-1106-0020) |
| **pGEX-KG** | pGEX-KG Expression plasmid containing a 5' GST tag. Ref. (63) |
| **pGEX-KG-$Ub_2$** | Herscovitch and Gilmore, unpublished |
| **pcDNA-FLAG C54A/C347A NEMO** | Ref. (27) |
| **Champion pET SUMO Protein Expression System** | Invitrogen, LifeSciences #K300-01 |
| **pET24b(+)-NEMO-1-419-WT** | Refs. (13, 28) |
| **pET24b(+)-NEMO-1-419-7XAla** | Refs. (13, 28) |

| | |
|---|---|
| **pET24b(+)-NEMO-1-419-9SG** | Ref. (28) |
| **SUMOstar-NEMO-44-195** | Ref. (13) |
| **SUMOstar-NEMO-110-195** | Ref. (13) |
| **pET-15b(+)-NEMO-44-419** | Genscript |
| **pET-28a(+)-NEMO-44-258** | Genscript |
| **pET-29(+)-NEMO-196-419** | TWIST Bioscience |
| **pET-29(+)-NEMO-OPTN-IVD** | TWIST Bioscience |
| **pET-29(+)-NEMO-OPTN-IVD** | TWIST Bioscience |
| **pET-SUMO-NEMO-252-419** | TA overhang PCR fragment containing amino acids 252-419 of human NEMO using primers 252-F and 419-R. Fragment was then subcloned into compatible TA linearized pET SUMO vector. |

| | |
|---|---|
| **pET-SUMO-NEMO-310-419** | TA overhang PCR fragment containing amino acids 310-419 of human NEMO using primers 310-F and 419-R. Fragment was then subcloned into compatible TA linearized pET SUMO vector. |
| **pET-SUMO-NEMO-344-419** | TA overhang PCR fragment containing amino acids 344-419 of human NEMO using primers 344-F and 419-R. Fragment was then subcloned into compatible TA linearized pET SUMO vector. |

**Supplemental Table 2. Primers used in this study.**

| Primer Name | Sequence |
|---|---|
| **252-F** | AGCGAACGTAAACGCGGTAT |
| **310-F** | GCCGATTTCCAGGCAGA |
| **344-F** | AAAGCAAGCTGTCAGGAATCT |
| **419-R** | TTATTCGATACATTCCATCACGTG |
